# OPTIMIZING PRE-PROCESSING OF NEAR INFRARED SPECTRA FOR PHENOMIC PREDICTION USING SINGULAR VALUE DECOMPOSITION

**DOI:** 10.64898/2026.05.10.724118

**Authors:** Clément Bienvenu, Jean-Michel Roger, Mamadou Séne, Sergio Antonio Castro-Pacheco, Mathilde Singer, Bakolinirina Laurencia Felaniaina, Nancy Terrier, Fabien De Bellis, David Pot, Hugues de Verdal, Vincent Segura

## Abstract

Phenomic prediction (PP) is a genetic value prediction method based on near infrared spectroscopy (NIRS). Spectra pre-processing is a key step in the analysis pipeline of PP and generally involves chemometrics methods. However, the choice of pre-processing is usually done either arbitrarily or through a search of the optimal set of methods and associated parameters. In this study, we propose to implement a singular value decomposition (SVD) step in the pre-processing pipeline where genetic values of spectra are estimated on a set of principal components instead of individual wavelengths. This way, estimations are based on a few informative, orthogonal and interpretable features of spectra instead of many correlated, uninformative wavelengths. We tested this pre-processing method on five datasets representing four plant species (maize, rice, sorghum and grapevine). Results show that estimating genetic values on components of raw spectra, that are not weighted by their eigenvalues, performs as well as doing it on spectra pre-processed with the best classical chemometrics methods in most cases, while requiring less parameter optimization. Moreover, this SVD step opens up possibilities for better understanding and selecting parts of the spectral information that are relevant for PP.

**Plain language summary:** Cultivated plants are the result of a breeding process during which their genetic values are used to select those to breed. Estimating these values requires heavy experimental means and is time consuming. Phenomic prediction is a low cost and high throughput method that is increasingly being used for this purpose. It often uses, as predictors, near infrared spectroscopy measurements that are easy to collect and thus routinely used in many species. However, near infrared spectra generally require pre-processing before being used in prediction. Currently used pre-processing methods arise from the chemometrics community, and still deserve a better in-depth appropriation by geneticists. In this study, we propose a pre-processing approach that performs as well as the best chemometrics pre-processing generally used, reduces computation time, and allows for a better understanding of what parts of spectral information are relevant for prediction.

**Core Ideas:**

- The SVD-based pre-processing performs as well as the best performing classical chemometrics pre-processing in most cases
- Using the SVD-based pre-processing reduces computing time of genetic value estimation and requires less parameter optimization than using classical chemometrics pre-processing
- Spectra are composed of chemical and physical information and classical pre-processing methods remove the physical part of the signal
- It is likely that chemical information is the most important for phenomic prediction even though physical information remains valuable
- Performance of the SVD-based pre-processing is likely due to a good estimation of the genetic part of spectra and the conservation of physical information of spectra

## INTRODUCTION

Phenomic prediction (PP) is a method for genetic value prediction inspired by genomic prediction (GP) (Rincent et al., 2018). PP works on a similar principle as GP but using high throughput phenotyping instead of molecular markers. In the case of the widely used H-BLUP model (phenomic equivalent to the G-BLUP model in GP (Van Raden, 2008)), the relationship matrix between individuals is calculated using high throughput phenotypic information instead of genomic based information. PP was originally developed using near infrared spectroscopy (NIRS) on plant tissues, and following studies have also tested hyperspectral, multispectral, RGB imaging or vegetation indices measured with drones, sometimes at several stages of the plants’ development. A growing body of evidence shows that PP using NIRS can reach a similar range of predictive abilities (PA) as GP (Adunola et al., 2024; Bienvenu et al., 2025; Brault et al., 2022a; De Salvio et al., 2024; de Verdal et al., 2024; Laurençon et al., 2024; Mora-Poblete et al., 2024; Rincent et al., 2018; Robert et al., 2022a; Robert et al., 2022b; Weiß et al., 2022). There are also few instances of PP performing better than GP (Gonçalves et al., 2021; Roscher-Ehrig et al., 2024; Zhu et al., 2021, 2022), and a more limited set of studies in which PP perform worse than GP (Cuevas et al., 2019; Jung et al., 2025; Zhu et al., 2022).

Some studies show that little information seems to be required to perform predictions compared to the amount of information usually collected. For example, Mróz et al. (2024) showed that only three wavelengths in the visible range acquired only once could predict yield of spring wheat with PAs ranging from 0.51 to 0.58. Also, Bienvenu et al. (2025) showed that randomly selecting less than 30 wavelengths could always reach the same PA as using all the wavelengths for seven traits in sorghum, and sometimes, even less than ten wavelengths were enough. These results stress out the fact that it is neither known which parts of spectra (or hyperspectral images) drive PP efficiency, nor what causes its PA to be rather high or low, in general and relatively to GP.

Spectra are generally pre-processed before being used in PP to isolate relevant information discriminating genotypes. The classical pre-processing pipeline uses chemometrics pre-processing methods (standard normal variate, detrend, first and second derivative, or combinations of those) followed by a model to extract the genetic part of the spectra. There is currently no way of knowing which chemometrics pre-processing will work the best on a given dataset without prior testing. Moreover, the choice of the best pre-processing can be tedious and require parameter optimization to maximize PA (Braun et al., 2025; Meyenberg et al., 2024). The modeling of the genetic part of spectra is then usually done after the pre-processing by calculating genotypic best linear unbiased estimations or predictions (BLUEs or BLUPs) of the absorbance or the reflectance at each wavelength in a linear mixed model. Then, these genotypic BLUEs/BLUPs are concatenated to form a new spectral matrix with one spectrum per genotype allowing the computation of a relationship matrix between genotypes.

Contrarily to geneticists, chemometricians have been using NIRS and other high-dimensional data for a long time as proxy for chemical composition of samples (Beć et al., 2020). They have developed a deeper understanding of spectral content and how to use it. We can thus learn from the chemometrics framework that spectra contain chemical and physical information (Chaminade et al., 1998; Fernandez Pierna et al., 2020). Physical effects are caused by the physical shape, size and structure of the measured sample that impact light scattering and absorbance according to the Kubelka-Munk theory (Kubelka & Munk, 1931). In classical chemometrics applications on solid samples, physical effects are generally linked to particle size of powders used for NIRS acquisition. This holds true for milled biological samples (e.g., flour, leaf powder, etc.). In the case of “intact biological material” (whole grain, leaf, wood, etc.), however, physical effects come from other sources. In the case of whole grains, they are most likely due to grain topography (size, curvature), texture of seed coat, non-uniformity of illumination and angle of presentation in the sample (Caporaso et al., 2018). Physical information is the main source of variance in spectra, whereas chemical information represents a minor part of spectral variance (Bull, 1991; Cowe & McNicol, 1985). Physical effects are visible in spectra as variation from the baseline (Chaminade et al., 1998; Liu et al., 2017), i.e., some spectra have consistently low value of absorbance while other have consistently high values (additive effects), and variance of absorbance values varies along wavelengths (multiplicative effects).

It is also known that wavelengths are generally correlated to one another (Cowe & McNicol, 1985). Thus, taken one by one, they are redundant and not very informative, which is not optimal to extract information from spectra such as their genetic values. Chemometricians, who try to extract chemical information from spectra generally work on spectral features rather than wavelengths to tackle this issue, using multivariate methods such as principal component analysis (PCA) or partial-least squares regression (PLSR), both methods being related to singular value decomposition (SVD).

Such chemometrics knowledge seems rather underexploited in the context of PP. Two elements can specifically be criticized and could be improved. Firstly, chemometrics pre-processing methods are meant to maximize spectral differences that are related to chemical differences by correcting scattering effects related to physical properties of samples (Caporaso et al., 2018; Chaminade et al., 1998). Removing physical information is justified in the classical application of NIRS where the goal is to predict chemical information such as protein or starch content. However, this may not be as relevant in the case of PP, as the physical information may be useful to discriminate genotypes. Secondly, the modeling of the genetic part of the spectra is done by computing hundreds, sometimes thousands of models on many correlated, noisy, and mostly uninformative wavelengths.

In this study, we propose to systematically add a SVD step in the pre-processing pipeline before the calculation of the genetic part of spectra. This way, principal components (PC) are used instead of wavelengths to estimate the genetic part of spectra. Replacing classical chemometric pre-processing by SVD allows to keep physical information in the spectra, and compute the genetic part of spectra on few independent informative features of spectra. Also, discarding the last PCs is known to be an efficient way to filter out noise (Antonelli et al., 2004), allowing a cleaner signal. This could in turn strongly reduce computation time when modeling the genetic part of spectra, as models can be fitted on a few dozens of PC instead of hundreds of wavelengths. Such approach relates to principal component regression (PCR), in the way that it reduces the dimensionality of spectra by representing it as a selected set of PC which are orthogonal and meaningful. However, it differs from PCR as, while PCR uses PC as explanatory variables to model response variables, the SVD-based pre-processing uses PC as response variables, to be modeled by explanatory variables (genotype, experimental design, environment).

Moreover, two variants of the SVD-based pre-processing were tested: either the genetic part of spectra was estimated on PC weighted by their eigenvalues (which are classical PCA components), or it was estimated on scaled PC. These two variants are motivated by the fact that the first PC of a PCA on spectra are more likely to bear the physical information of spectra as this is the information that causes the most variability in spectra (Bull, 1991; Cowe et al., 1985). Thus, working on PC weighted by their eigenvalues is likely to emphasize the role of physical information in prediction, since the first PC will have a greater impact on the final SVD-pre-processed spectra. On the other hand, working on scaled PC is likely to emphasize the role of both physical and chemical information in spectra as all PC will have the same impact on the final SVD-pre-processed spectra. Comparing these two variants to classically pre-processed spectra, which theoretically only retain chemical information could help understand which part of the NIRS signal is the most valuable for prediction.

In the present study, we tested and compared the predictive abilities achieved with classical chemometrics pre-processing methods, and SVD-based pre-processing methods (weighted and not weighted), with or without combination to classical chemometrics pre-processing methods. This was done on five datasets, three of which had previously been used in phenomic prediction studies (Brault et al., 2022a; De Salvio et al., 2024; de Verdal et al., 2024; Lane et al., 2020). These datasets encompass three population types (half-diallel, diversity panel, breeding material), six types of NIRS acquisition material (NAM) (grain, leaf, straw, wood, flour, ground straw), six types of traits (yield, phenology, fruit/grain weight, vigour and spike/ear/grape morphology) on four species: rice (*Oryza sativa*), maize (*Zea mays subsp. mays*), grapevine (*Vitis vinifera*), and sorghum (*Sorghum bicolor*).

## MATERIAL AND METHODS

### Datasets

Five datasets from four different plant species were used in this study: rice, maize, grapevine, and sorghum. They correspond to three types of populations (breeding material, diversity panel, and a half diallel crossing scheme) and include six types of traits representing yield, grain/fruit weight (GW), plant height (PH), phenology (PHENO), morphology of the fruit/grain bearing organ (panicle, ear, spike, bunch of grapes, MORPH), and vigour (VIG). Three datasets have already been used for PP analyses (Brault et al., 2022a; De Salvio et al., 2024; de Verdal et al., 2024; Lane et al., 2020). Such previously used datasets will be described briefly to provide readers with essential information for the understanding of this study. Readers may refer to cited studies for complete descriptions of the datasets.

#### Rice dataset 1

This dataset was used for PP by de Verdal et al. (2024). Details on field experiment, hyper-spectral measurements, and genotyping can be found in (Rakotomalala et al., 2021; Rakotoson et al., 2021, 2017).

A panel of 190 accessions representing the working collection of the upland rice breeding program conducted in Madagascar by FOFIFA (Malagasy National Research Center for Rural Development) and CIRAD (French Agricultural Research Center for International Development) was used. These accessions were grown during two consecutive seasons (2014-2015 and 2015-2016) in split-block designs with two replications in an alpha lattice lay out. Each replication consisted of 14 blocks, further subdivided into two sub-blocks for evaluating nitrogen fertilization levels (low and high).

The traits used were plant height (PH), grain yield (YIELD), thousand grain weight (GW), days to flowering (PHENO), number of panicles per square meter (VIG). NIR reflectance spectra were collected on harvested grains milled to 1 mm. Spectra were acquired for each plot (each accession in each sub-block) and include data from 1,500 wavelengths ranging between 1,000 and 2,500 nm with a 1 nm step. Four technical repetitions were averaged to obtain the spectra of each plot.

#### Rice dataset 2

This dataset has never been used in any previous work. It comprises 350 breeding lines (F_5_ and F_6_ lines) from the FOFIFA–CIRAD upland rice breeding program, evaluated during the 2024–2025 growing season across two experimental sites in Madagascar: the FOFIFA Andranomanelatra Research Station (19°47’20.76"S, 47°06’41.76"E) and Talata Andraikiba (19°52’ 12.00"S, 46°59’02.40"E). At the Andranomanelatra site, trials were implemented under two contrasting fertility conditions (high and low fertility), whereas at Talata Andraikiba, experiments were conducted exclusively under high fertility. Altogether, these conditions defined three distinct environments for subsequent analyses.

In each environment (location/management combination), an augmented p-rep design (20% of replication per environment) was used. Blocks consisted of eight plots and included one check genotype (FOFIFA 186), which was repeated in every block, in addition to the partially replicated 350 genotypes. Experimental plots measured 1 x 2.4 m. at the Andranomanelatra site, and 1 x 2 m. at Talata Andraikiba. Sowing hills were spaced at 20 × 20 cm, with five seeds manually sown per hill. Sowing was carried out between 20 and 25 October 2024, depending on the environment. All environments received 500 kg/ha of dolomite and 5 t/ha of manure before sowing. In addition, high-fertility plots received 80 kg/ha of NPK fertilizer (11–22–16) at sowing and 60 kg ha^−1^ of urea before flowering.

The measured traits were plant height (mean of 5 plants per plot, PH), weight of harvested grain at each plot (YIELD), weight of 1000 grains (GW), flowering date (PHENO), panicle length (mean of 5 plants per plot, MORPH), and number of panicles per hill (mean of 5 plants per plot, VIG).

NIR reflectance spectra were measured on whole grains and ground leaves after harvest. Spectral acquisition was performed using an ASD LabSpec 4 spectrometer, covering 1,500 wavelengths in the range of 1,000–2,500 nm.

#### Maize dataset

This dataset was first reported by Farfan et al. (2015) and further used for PP in Lane et al. (2020) and De Salvio et al. (2024)..

A set of 346 diverse F1 hybrids were grown in 2011 and 2012 with two watering conditions (well-watered and water stress) in each year, in two replicates in each year/watering condition combination, with a randomized complete block design (RCBD) at the Texas A&M AgriLife Research Farm.

Five traits were used, namely plant height (PH), grain yield (YIELD), 500-kernel weight (GW), days to silking (PHENO), and ear height (MORPH).

NIR reflectance spectra were collected on harvested whole kernels, and consisted of every wavelength from 4,000 to 10,000 cm^-1^, which corresponds to an interval between 1,000 and 2,500 nm. Spectra were acquired for each plot (each hybrid in each year/watering condition/repetition).

#### Grapevine dataset

This dataset was used for PP in Brault et al. (2022a). It comprises two distinct populations for which analyses were conducted separately.

The first population is a diversity panel composed of 279 accessions grown at the Domaine du Chapitre experimental vineyard of Institut Agro, Montpellier, in Villeneuve-lès-Maguelone (France). The panel is replicated in five complete blocks (RCBD), each accession being represented by one plot of a single vine in each block. The second population is a half-diallel population composed of 676 genotypes, grown in the same location. Genotypes are derived from ten bi-parental crosses between five varieties. The population is replicated in 2 complete blocks (RCBD), each genotype being represented by one plot of two consecutive vines in each block.

Phenotypic data were collected in 2011 and 2012 for the diversity panel and between 2013 and 2017 for the half-diallel yielding two years of measurement for each trait in each population. For both populations, traits used were the number of clusters per plant (YIELD), mean berry weight (GW), véraison date (PHENO), bunch compactness (MORPH), and vigour (VIG).

NIR reflectance spectra were measured in both populations on dried and milled wood and dried adult leaf disks collected during two consecutive years (2020 and 2021). Wavelengths ranged from 350 to 2,500 nm, with a 1 nm step.

#### Sorghum dataset

This dataset was initially designed for an association mapping study of grain quality and has never been used for PP. The population developed in this study comprises 303 accessions representing a wide range of ecotypes, plant architectures and grain quality traits, covering the five main cultivated races (Caudatum, Durra, Bicolor, Kafir, and Guinea) and ten intermediate races. The geographical origin of the accessions also encompassed the major centers of sorghum diversity in Africa and Asia, as well as cultivated lines from the Americas, India, and the Mediterranean region. It consists of 30 accessions from the CIRAD Core Collection (Deu et al., 2006), 103 accessions from the GCP Reference Set (Billot et al., 2013), 119 accessions from the Sorghum Association Panel (Sukumaran et al., 2012; Luo et al., 2016; Cuevas et al., 2018; Boatwright et al., 2022), and 51 accessions selected from the literature based on criteria related to grain quality, particularly protein and starch contents as well as protein digestibility (Axtell et al., 1981; Hess et al., 2002; Yan et al., 2011; Duressa et al., 2020 ; Kardes et al., 2021).

The 303 accessions of the panel were sown in mid-May 2023 and mid-May 2024 at Rivières (Southwest France: 43°54’43.60"N, 1°59’22.50"E) and Mondonville (Southwest France: 43°40’29.64"N, 1°17’34.04"E), under rainfed. At each site, an augmented block design was established. The tested accessions were arranged by plant height to avoid shading between plants, without replication, while four check genotypes were replicated across blocks, to account for field heterogeneity in the statistical analyses. Each accession was sown in a plot consisting of two rows spaced 0.8 m apart. Each row was 5 m long and contained 64 plants, corresponding to a density of 160,000 plants/ha. Standard local agronomic practices were applied to ensure optimal plant growth and development, including supplementary irrigation, fertilization, and pest management. Due to differences in earliness among accessions, harvests were carried out between September and October at both sites and in both years. At physiological maturity, five panicles were randomly sampled from each plot. After harvest, panicles were dried and threshed mechanically, and remaining glumes were removed as much as possible.

Traits used for PP were plant height (PH), thousand kernel weight (GW), and days to flowering (PHENO).

NIR reflectance spectra were collected on whole grains, or grains milled to a 1 µm powder. In both cases, spectra were acquired with a TANGO spectrometer (Bruker Optik GmbH, Ettlingen, Germany). For whole grains, four spectra were acquired and averaged for each sample, while two spectra were acquired and averaged for milled grains. Spectra consist of wavelengths between 11528 and 3944 cm^−1^ (867 and 2535 nm) with 8 cm^−1^ steps. Spectra were only acquired in one environment (Rivière 2024).

### Estimation of genotypic BLUEs

For each dataset, the genotypic BLUEs estimation models were built upon the ones published in their respective articles. Some adjustments were made to have genotypic values always estimated with fixed genetic effects as recommended in Holland & Piepho, (2024), which was not the case in all original models.

#### Rice dataset 1

As in de Verdal et al. (2024), genotypic BLUEs were estimated within environments (year/nitrogen treatment combination) with the following model:

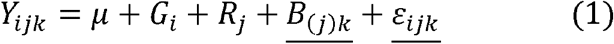

Where *Y*_*ijk*_ *i*s the observation of the genotype *i* in block *k* of repetition *j*.*G*_*i*_ *i*s the fixed effect of genotype *i, R*_*j*_ is the fixed effect of repetition *j*,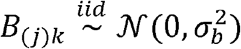 is the random effect of block *k* nested into repetition *j*, 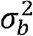 being the block variance, .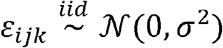 is the random residual, σ^2^ being the residual variance.

Environment specific genotypic BLUEs were estimated with the main effect of genotypes within environments.

#### Rice dataset 2

As this dataset has never been used, no genotypic value estimation model was available. We fitted a model to account for the effects of the experimental design. Environments were not a variable of interest but were fitted as a fixed effect because there are only three environments. Blocks and the interactions between genotypes and environments were not variables of interest and were thus fitted as random effects:

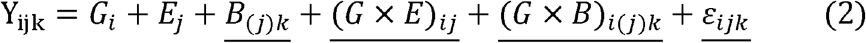

Where *Y*_*ijk*_ *i*s the observation on genotype *i* in block *k* of environment *j, G*_*i*_ *i*s the fixed effect of genotype *i, E*_*j*_ *i*s the fixed effect of environment *j* as a combination between location and management condition,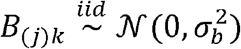 is the random effect of block *k* nested into 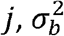 environment *j*, being its associated variance,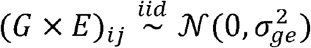 is the random effect of the interaction between genotype *i* and environment 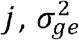 being its associated variance,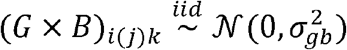 is the random effect of the interaction between genotype *i* and block 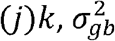, being its associated variance, .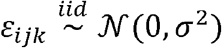 is the random residual, σ^2^being the residual variance. Across environments genotypic BLUEs were estimated as the G effect.

#### Maize dataset

As in Lane et al. (2020), genotypic BLUEs were estimated within environments. In this article, a model with a genotypic and a block effect was computed. However, no block information could be found in the available dataset from this article. Instead, a row, a range, and a repetition column were available. In De Salvio et al. (2024), which uses the same dataset, genotypic adjusted means were computed in a model using the row, range and repetition information. Therefore, the following model was computed in each environment:

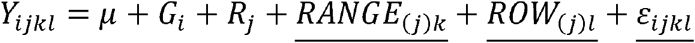

Where *Y*_*ijkL*_ *i*s the observation of genotype *i* in range *k* and row *l* of repetition *j*.*G*_*i*_ *i*s the fixed effect of genotype *i, R*_*j*_ is the fixed effect of repetition,,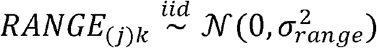 is the random effect of range nested into repetition, 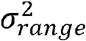 being the range variance,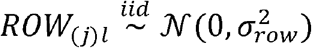 is the random effect of row nested into repetition, 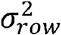 being the row variance, 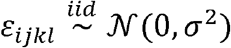 is the random residual, σ^2^being the residual variance. In De Salvio et al. (2024), the effect of repetition was considered random as the range and row effects. We computed it as fixed here because there are only two repetition per environment. Environment specific genotypic BLUEs were estimated as the G main effect within environments.

#### Grapevine dataset

Models used to estimate the genotypic adjusted mean of phenotypic data can be found in Flutre et al. (2022) for the diversity panel, and in Brault et al. (2022b) for the half diallel population. The model used to estimate the genotypic adjusted means of spectra is the same for both the diversity panel and the half diallel population and is described in Brault et al. (2022a). However, the models used to estimate the genotypic adjusted means of the phenotypes in the half diallel and the diversity panel were not the same, despite similar experimental designs and similar genetic structures (genotypes nested into crosses for the half diallel, and nested into gene pool for the diversity panel). Likewise, the models used to estimate the genotypic adjusted means of the spectra were not the same as the ones used for the phenotypes, even though spectra are phenotypes. Thus, we decided to use a model built upon the original spectra genotypic adjusted mean estimation model from Brault et al. (2022a), and added some effects present in the phenotypic genotypic adjusted mean estimation model in Brault et al. (2022b). The final model used in our study is the following:

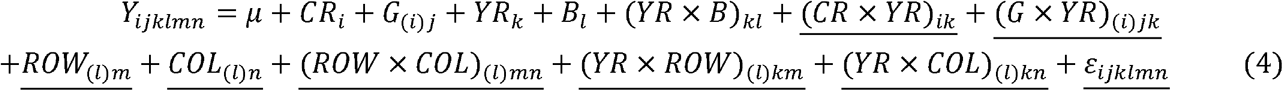

Where *Y*_*ijklmn*_ *i*s the observation of genotype *j* from cross (or gene pool) *i* in year *k*, block *l*, at line *m* and column *n* of the field layout, *CR* _*i*_ *i*s the fixed effect of cross *i* for the diallel population, and the fixed effect of gene pool *i* for the diversity panel, *G*_(*i*)*j*_ *i*s the fixed effect of genotype *j* nested into cross/gene pool *i, YR*_*k*_ *i*s the fixed effect of year *k, Bi* is the fixed effect of block *l*,(YR ×*B*)_*kl*_ *i*s the fixed effect of the interaction between block *l* and year *k*,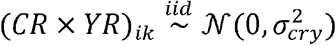 is the random effect of the interaction between cross *i* and year *k*, 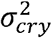 being its associated variance, 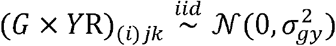 is the random effect of the interaction between genotypes (*i*)*j* and year 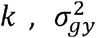 being its associated variance,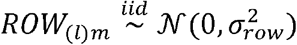 is the random effect of field row *m* nested into block 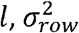 being the row variance, 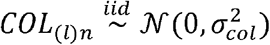 is the random effect of field column *n* nested into block 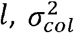 being the column variance, 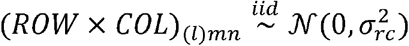 is the random effect of the interaction between row *m* and column 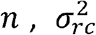 being its associated variance, 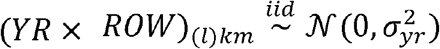 is the random effect of the interaction between year *k* and row *m*,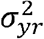 being its associated variance, 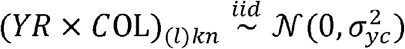 is the random effect of the interaction between year *k* and column 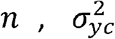 being its associated variance,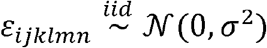 is the random residual, σ^2^being the residual variance.

Cross and genotype effects were fitted as random in Brault et al. (2022a), which is not the case here where these effects were considered fixed. Interactions between row/column and year were present in Brault et al. (2022b) for phenotype genotypic BLUEs estimation, but not in Brault et al. (2022a) for spectra genotypic BLUEs estimation. Across year/environment genotypic BLUEs were estimated as the sum of the cross/gene pool and genetic main effects.

#### Sorghum dataset

As for the second rice dataset, the sorghum dataset has never been used in any previous work. We thus fitted the following model to account for the effects of environments of experimentation and of the experimental designs. There were only two environments and two years so these effects and their interaction were considered fixed. Blocks and the interactions between genotypes locations and years were not variables of interest and were considered random:

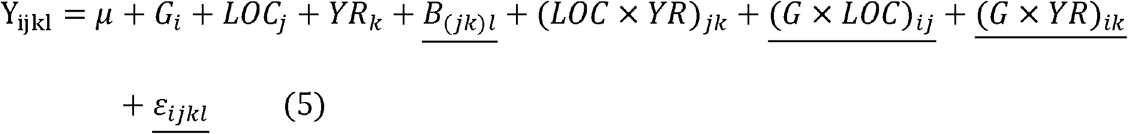

Where *Y*_*ijk*_ *i*s the observation on genotype *i* in block *l* of environment *j* and year *k, G*_*i*_ *i*s the fixed effect of genotype *i, L O C*_*j*_ *i*s the fixed effect of location *j, YR*_*k*_ *i*s the fixed effect of year *k*,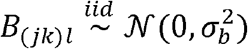 is the random effect of block *l* nested into location *j* and year 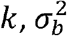 being its associated variance, (*L O C* ×*YR*)_*jk*_ *i*s the fixed effect of the interaction between location *j* and year 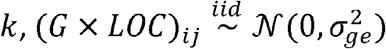 is the random effect of the interaction between genotype *I* and location 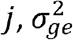 being its associated variance, 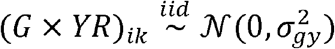 is the random effect of the interaction between genotype *i* and year 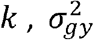 being its associated variance, 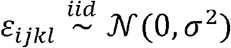 is the random residual, σ^2^ being the residual variance. Across environment Genotypic BLUEs were estimated as the G effect. In this dataset, days to flowering (PHENO) was measured in only one environment. Thus, the *L O C* effect and its interactions were removed in the model to estimate the genetic BLUEs for this trait. The lack of repetitions of many genotypes and their field arrangement by plant height also creates limitations for a good estimation of genotypic BLUEs, and may have introduced confounding between genotypes and field heterogeneity.

### Analysis of variance and heritability

Such analyses have previously been done for the rice dataset 1, the maize dataset and the grapevine dataset in their respective articles. For the rice dataset 2 and sorghum datasets, the models used to estimate variance components were the same as the ones used for genotypic BLUEs estimation but with all terms considered random (model 2 for rice and 5 for sorghum).

As these datasets are p-rep and augmented design, respectively, the heritability was calculated according to Cullis et al. (2006):

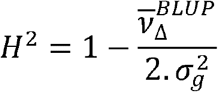

Where 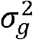 is the genetic variance and 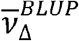 is the average standard error of the genotypic BLUPs.

### Spectra pre-processing

Four classical pre-processing methods were used on raw spectra: normalization with the standard normal variate method (snv), detrend (dt), first order derivative after snv (d1), and second order derivative after snv (d2). All those pre-processing methods were used after a smoothing of raw spectra using the Savitsky-Golay algorithm (filter length of 21 wavelengths, polynomial order of 1, and derivation order of 0). Derivative were also done with the Savitsky-Golay algorithm. For d1, parameters were: filter length of 41 wavelengths, polynomial order of 2, and derivation order of 1. For d2, parameters were: filter length of 81 wavelengths,polynomial order of 3, and derivation order of 2.After this pre-processing step, the models previously presented for the estimation of genotypic BLUEs were fitted on spectra to estimate the genotypic BLUEs of the reflectance at each wavelength. This step allows to have one spectrum per genotype (or per genotype-environment combination for rice 1 and maize datasets) to calculate relationship matrices between genotypes using spectral information. An exception was made for the sorghum dataset for which spectra were acquired in one environment and one year only, so we used model 5 with the effects of genotypes and blocks only.

Lastly, two SVD based pre-processing were tested. First, a SVD was made on centered but not scaled raw spectra (as wavelength are all in the same unit), decomposing the spectra as *UDV′* matrices. Then, two ways of calculating the genotypic adjusted means were used: either on the columns of the *U* matrix (later called the U method), or on the columns of the *UD* matrix which corresponds to components of a PCA (later called the UD method). Let *U* and (*UD*) _*g*_ be the genotypic adjusted means of *U*_*g*_ and *UD*_*g*_ respectively. Genotypic adjusted means of components were back transformed into genotypic adjusted means of wavelengths by calculating *U*_*g*_*V′* for the U method and (*UD*) _*g*_*V′* for the UD method. Different number of components were tested for the back transformation to spectra: from one to 10 components by steps of one, and from 10 to 100 components by steps of five.

The difference between the U and the UD methods is that in the U method, all components contribute equally to the estimation of the genotypic adjusted means, while in the UD method, components are weighted by their eigenvalues (the part of spectral variance that they represent). In other words, the U methods gives a greater importance to components representing a small part of variance than the UD method. The UD method is equivalent to working on components of a PCA, and the U method is equivalent to working on scaled components of a PCA.

### Prediction models

The model used for prediction was:

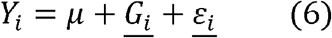

Where *Y* is the vector of genotypic values estimated with models described in section “Estimation of genotypic BLUEs”, 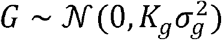 is the random effect of genotypes, 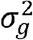 being the genetic variance, and *K* being the relationship matrix calculated with spectra, and ε ∼ *N*(0, *Iσ*^2^) the random residuals, being the identity matrix, and *σ*^2^ the residual variance. The relationship matrix was calculated as 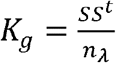, *S* being the scaled and centered matrix of spectra genotypic BLUEs, and *n*_*λ*_ the number of columns of *S*, i.e., the number of wavelengths composing the spectra. For the maize and rice dataset 1, predictions models were computed within each environment.

### Cross validation and predictive ability

A nested cross validation scheme was implemented for each dataset-NAM combination, where 70% of genotypes were used as a training set, and 30% as a validation set, constituting the outer loop of the scheme. Among the training set, an inner five-fold cross validation was implemented to test the performance of each classical chemometrics pre-processing method, and different number of components for the SVD-based methods.

For the classical chemometrics methods, spectra were pre-processed, then the genotypic adjusted means of wavelengths were computed using all genotypes. All pre-processing methods were tested in the inner loop, and only the best performing method was kept to be used in the outer loop.

For the SVD-based methods, SVD and genotypic adjusted means were computed only on the training set. The back transformations to spectra using different number of components were all tested in the inner loop, and only the best performing number of components was used in the outer loop. To perform the outer loop, spectra from the validation set of the outer loop were projected in the space of components. Genotypic adjusted means were then re-computed with all genotypes and the prediction of new genotypes was done. Let *S*_*val*_ be the spectra kept for the outer validation set. The projection of *S*_*val*_ onto the space of components was done by calculating *S*_*val*_ *VD*^-1^ for the U method and *S*_*val*_ *V* for the D method.

For each trait in each data-set-NAM combination, 100 repetitions of the whole cross validation scheme were done.

Figure 1 is a schematic representation of the pre-processing methods and cross validation schemes implemented.

**Figure 1:**
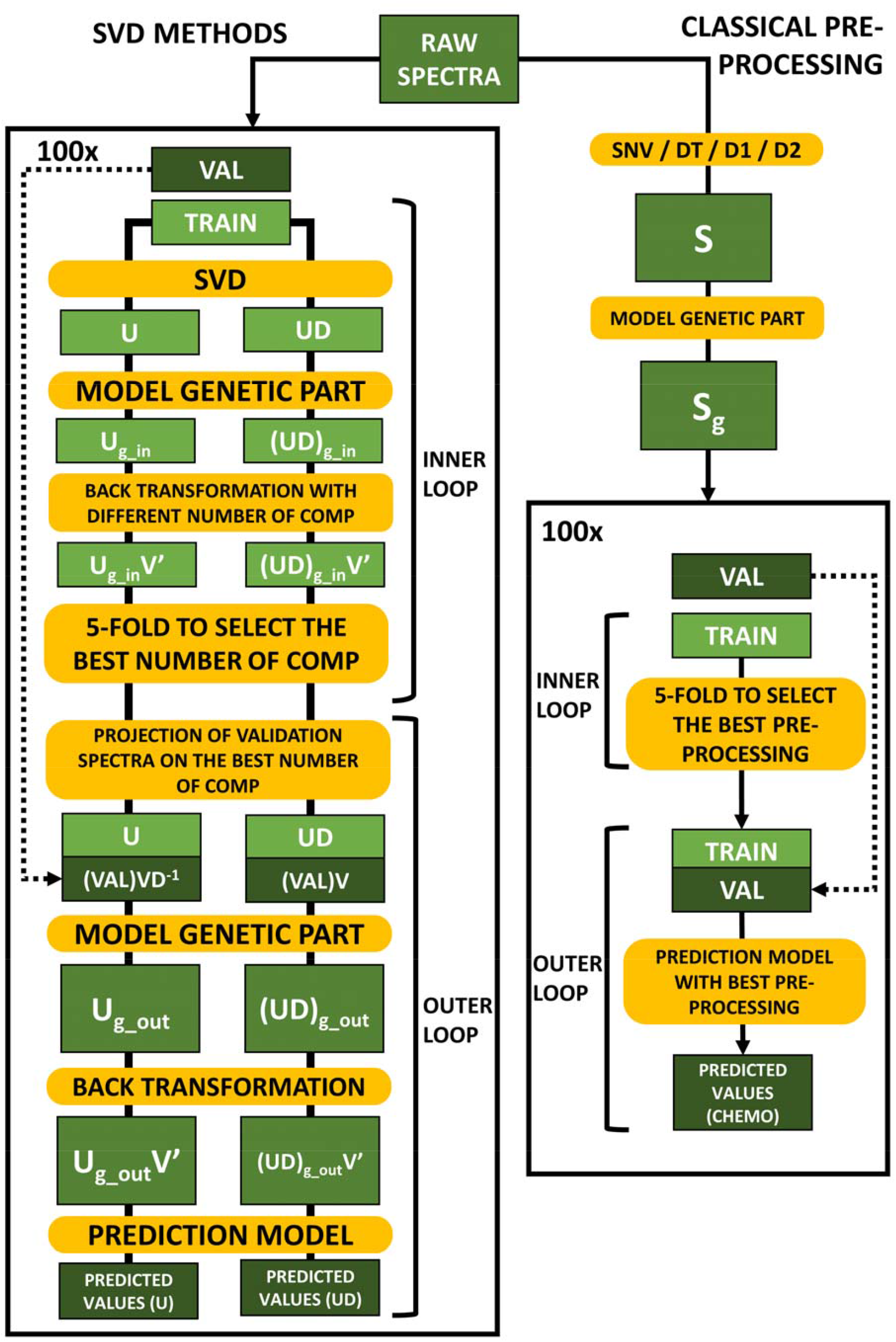
Representation of the pre-processing methods and cross validation scheme. SVD = singular value decomposition, TRAIN = outer training set on which a 5-fold cross validation is applied in the inner loop, VAL = outer validation set, S = pre-processed spectra, S_g_ genetic BLUEs of wavelengths of pre-processed spectra, snv = standard normal variate, dt = detrend, d1 = first derivative after snv, d2 = second derivative after snv, U, D and V = matrices obtained by the SVD decomposition on TRAIN, U_g_in_ / (UD)_g_in_ = genetic BLUEs of the columns of U or of UD, U_g_out_ / (UD)_g_out_ = genetic BLUEs of the columns of U or of UD in the outer loop after projection of the validation set in the space of U.

**Figure 2:**
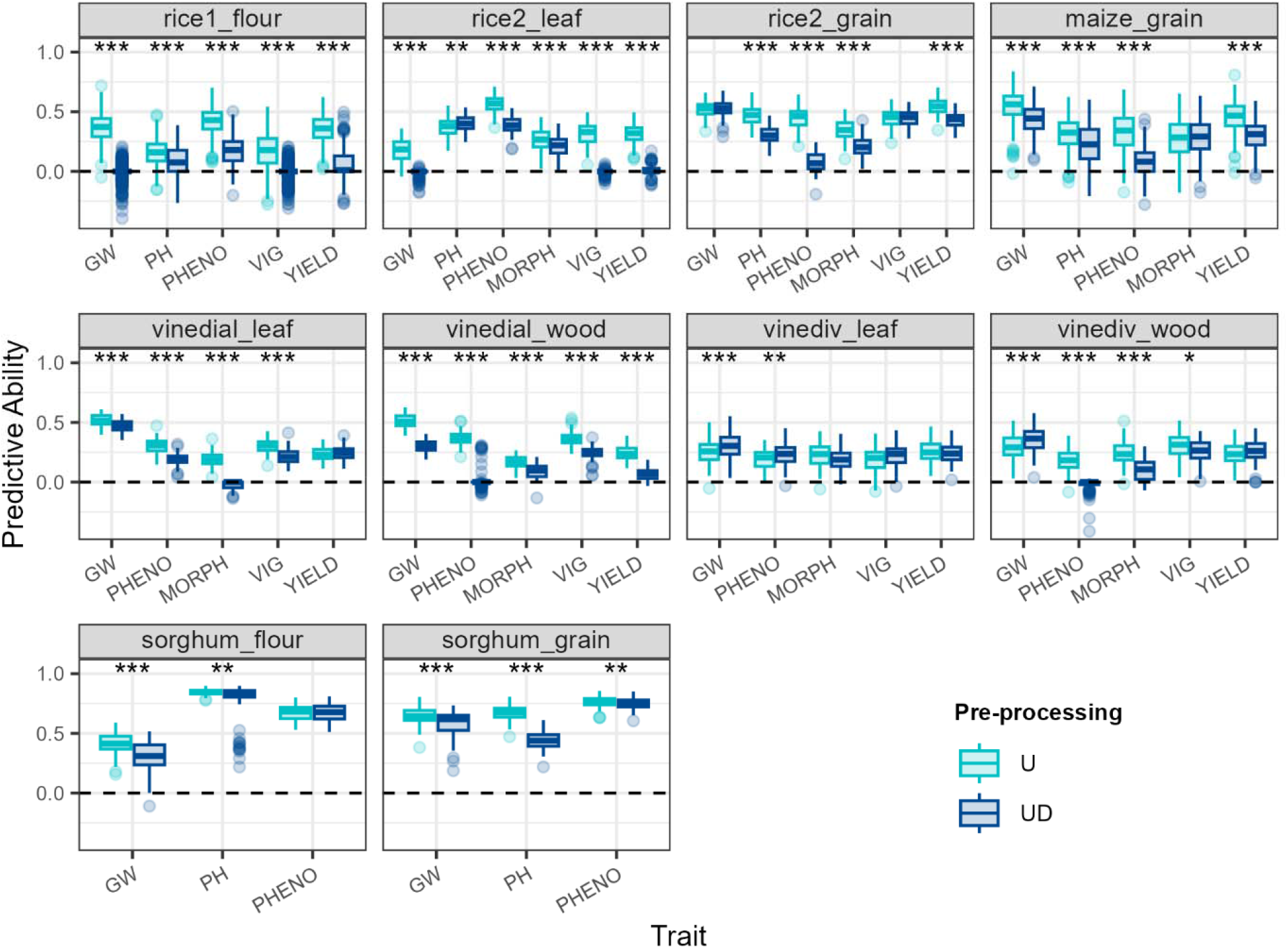
Comparison of predictive abilities between the U and UD methods. In the U method, genotypic BLUEs were estimated on the U matrix of a singular value decomposition of raw spectra. In the UD method, genotypic BLUEs were estimated on the UD matrix of a singular value decomposition of raw spectra. For each cross-validation repetition, trait, NIRS acquisition material (NAM) in each dataset, the number of components kept was determined by an inner five-fold cross validation on the training data. GW = grain weight or berry weight, PH = plant height, PHENO = phenology, MORPH = morphology of the grain/fruit bearing organ, VIG = vigour. Stars indicate significative differences between U and UD methods for a given trait, a given NAM in a given dataset. Differences were tested with a paired Wilcoxon test and p-values were adjusted with the Bonferroni method across data sets NAM and traits. * = p-value < 0.05, ** = p-value < 0.01, *** = p-value < 0.001.

### Predictive abilities and statistical tests

Predictive abilities were computed as the spearman correlation between predicted values of the outer validation loop and true genotypic BLUEs. Comparisons between pre-processing methods were always made between two methods using paired Wilcoxon tests for each data-set-NAM-trait combination. As a non-parametric method, the Wilcoxon test does not require normally distributed data and is suited for correlations that are bounded between -1 and 1. Tests were made paired to account for the fact that models were evaluated on the same sets of cross validation. As multiple tests were made, a Bonferroni correction was applied on p-values. For example, to compare the U and UD methods, one paired test was done for each trait in each dataset for each NAM, and the Bonferroni correction was applied on the p-values of all tests across datasets, NAM and traits.

### Software and packages

All analyses were conducted using the R software (R Core Team, 2025). Genotypic value estimations were done using the lme4 package (Bates et al., 2015), heritability calculation was done using the agriutilities package (Aparicio & Bornhorst, 2026), prediction models were fitted using the sommer package (Covarrubias-Pazaran, 2016), and graphs were made with the ggplot2 package (Wickham, 2016). The tidyverse environment was used all along the analyses for data management (Wickham et al., 2019).

## RESULTS

### Phenotypic data analysis

Readers should refer to de Verdal et al. (2024) for rice dataset 1, to De Salvio et al. (2024) and Lane et al. (2020) for the maize dataset, and to Brault et al. (2022b) and Flutre et al. (2022) for the grapevine dataset.

Summary statistics for traits of the sorghum and rice dataset 2 are available in table S1. For both datasets, PHENO and GW were the traits with the least variation across environment. PH had higher but moderate variations across environments for both datasets.

For the rice dataset 2, VIG mean and standard deviation were considerably higher in the TAL environment compared to the AND-FM and AND-FU environments. On the other hand, MORPH means had low variations between environment. The AND-FU environment had considerably lower yield than AND-FM and TAL environments.

Partitions of variance are available in figure S1 and S2 for the sorghum and rice dataset 2 respectively. For the sorghum dataset, partitions of variance showed that genetic variance was the major variance component, with heritability of 0.83 for GW, 0.96 for PH and 0.89 for PHENO. It is also striking that block variance was null for all traits. However, in this dataset, there were confounding effects between genotypes and field heterogeneity. Thus, it is likely that the estimated genetic variance was inflated, because it includes both genetic and block variances. This also means that genotypic BLUEs were likely poorly estimated.

For the rice dataset 2, residual variance was the highest for GW and MORPH, while environmental variance was the main variance component for PH, PHENO, VIG and YIELD. Genetic variances were low, always under 25% of total phenotypic variance, with heritability of 0.48 for GW, 0.61 for MORPH, 0.74 for PH, 0.67 for PHENO, 0.16 for VIG, and 0.52 for YIELD.

### Comparison between U and UD methods

Figure 1 represents the comparison between the two variants of the SVD-based pre-processing method: either genotypic BLUEs were estimated on scaled components (U method) or on classical PCA components (UD method). SVD was performed on raw spectra. Across all traits and datasets, the U method significantly outperformed the UD method 35 times (73%), the two methods were equivalent nine times (19%), and UD outperformed U four times (8%). It is noticeable that the vinediv dataset with leaf spectra had contrasting results with two occurrences of UD outperforming U, and two occurrences of U and UD having similar performance. Otherwise, the U method almost consistently outperformed the UD method, and no clear pattern emerged to explain the cases where it did not. These results showed that when adding a SVD step to estimate the genotypic BLUEs of raw spectra, working on the U matrix (or scaled components of a PCA) was more effective than working on the UD matrix (or PCA components). Thus, the comparison between classical pre-processing and SVD-based pre-processing was further made by using only the U method.

### Comparison between classical pre-processing and SVD methods

Figure 3 represents the comparison between the U method and the best classical pre-processing method. In each repetition of the cross validation in each trait, data set and NAM combination, the choice of the best pre-processing method was made by testing all four classical pre-processing methods in an inner five-fold cross validation on training data only, retaining the one having the best predictive ability to test on the validation data and produce the final predictive ability shown in the figure. The same method was used to choose the number of components to be kept with the U method.

**Figure 3:**
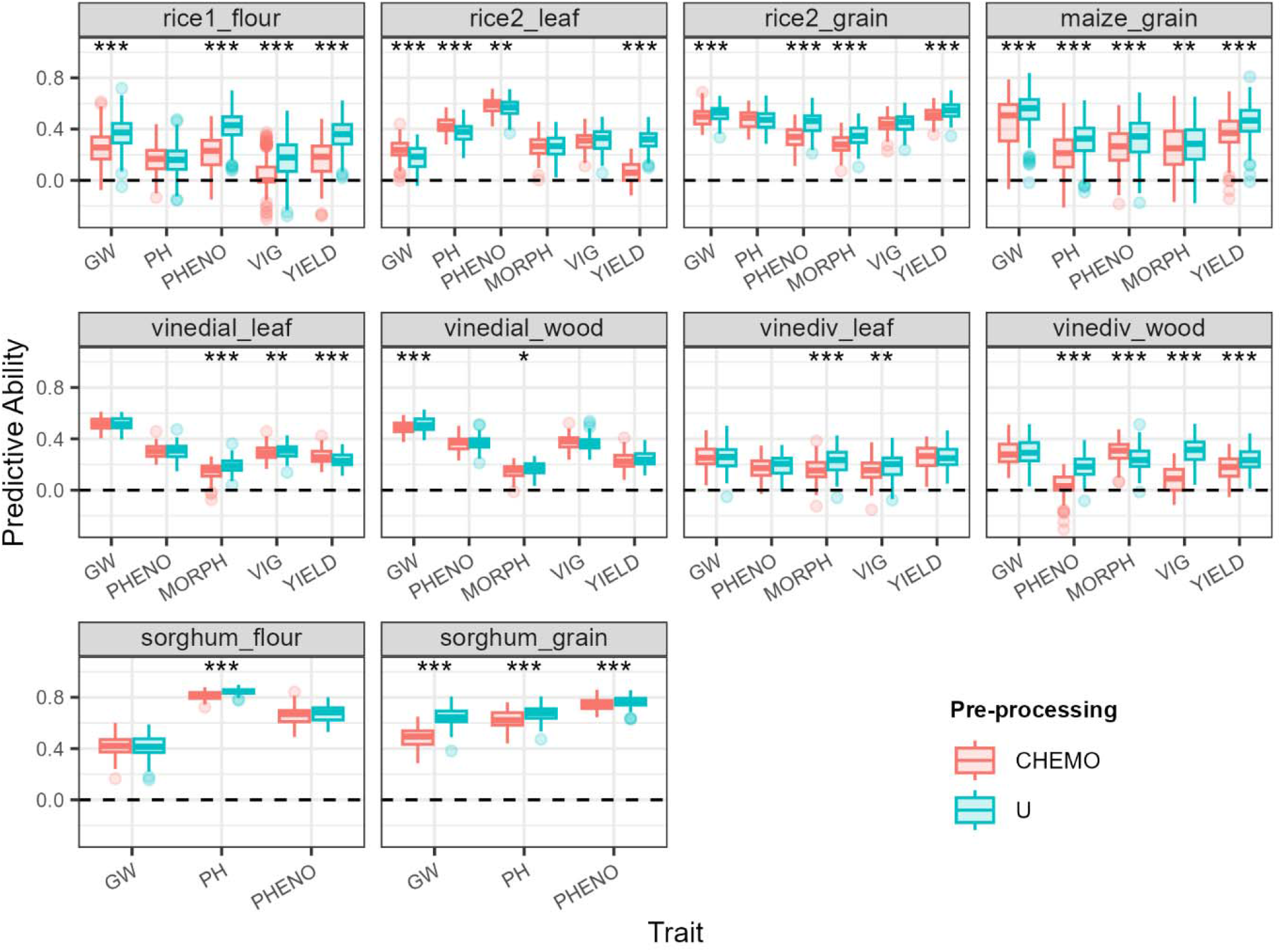
Comparison of predictive abilities between the U and classical pre-processing methods. In the U method, genotypic BLUEs were estimated on the U matrix of a singular value decomposition of raw spectra. In the CHEMO method, genotypic BLUEs were estimated on the wavelengths of pre-processed spectra. For each cross-validation repetition, trait, NIRS acquisition material (NAM) in each data set, the number of components kept and the best classical pre-processing method were determined by an inner five-fold cross validation on the training data. GW = grain weight or berry weight, PH = plant height, PHENO = phenology, MORPH = morphology of the grain/fruit bearing organ, VIG = vigour. Stars indicate significative differences between U and CHEMO methods for a given trait, a given NAM in a given data set. Differences were tested with a paired Wilcoxon test and p-values were adjusted with the Bonferroni method across data sets NAM and traits. * = p-value < 0.05, ** = p-value < 0.01, *** = p-value < 0.001.

Results show that the U method significantly outperformed the best classical pre-processing 27 times (56%), both methods were equivalent 16 times (33%), and the best classical pre-processing significantly outperformed the U method five times (10%). There was no trait or NAM for which results were strongly consistent across data sets, and no data set for which results were consistent across traits or NAM.

Each significance test was performed by comparing two paired samples of 100 repetitions and significant differences were generally small when replaced in the context of predictive ability gain. Figure 4 represents the histogram of differences between the U and best classical pre-processing methods for each repetition of cross validation in each data-set-NAM-trait combination (calculated as U-CHEMO). The distribution was skewed toward positive differences indicating that the U method had more chances of performing better than worse compared to classical pre-processing methods. Still, the first and third quartiles of the distribution were - 0.016 and 0.136, meaning that half of the gains of U over classical pre-processing were between these values. Thus, results showed that the U method generally performed as well as the best classical pre-processing methods with more chances of performing better than worse.

**Figure 4:**
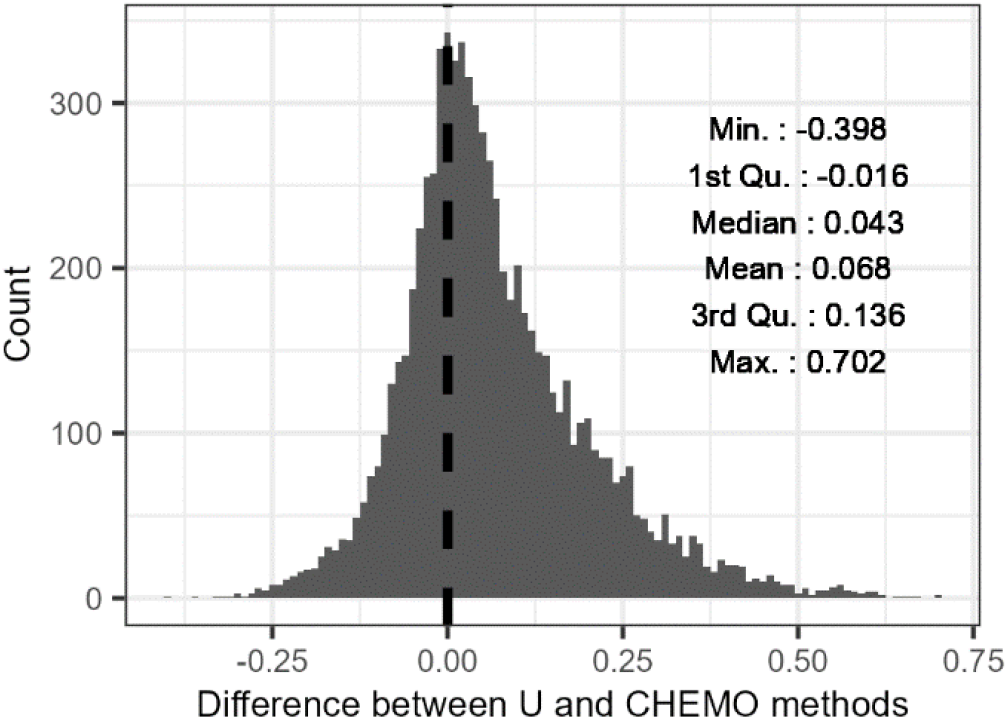
Histogram of the differences in predictive ability between the U method and the best classical pre-processing method. Differences were computed for each repetition of cross validation, for each trait in each data-set-NAM combination. Positive values indicate that the U method performed better than the best classical pre-processing, negative values indicate the opposite. Each bar represents an interval of 0.01.

Figure S3 is similar to figure 3 but adds the results of raw spectra (no pre-processing at all) and the best combination of classical and the U method (classical pre-processing being made before the U method, and choice of classical pre-processing and number of PC made with an inner 5-fold cross validation on training data only). This figure shows that PA achieved with raw spectra were generally lower and that combining classical and SVD-based pre-processing generally performed as well as the U method on raw spectra.

### Optimization of classical pre-processing and number of components

Figure 5 represents the number of times each classical pre-processing was the best performing in each data-set-NAM combination across traits (according to the inner loop of cross validation). Results show that the most performant classical pre-processing methods were generally d1 or d2 which were combinations of snv and a derivative of order one or two. Indeed, for all data-set-NAM combination except sorghum_flour, either d1 or d2 was the best performing method in most cases. This was especially true for vinedial_leaf, vinedial_wood, vinediv_leaf and sorghum_grain where d1 or d2 were the best performing in 94%, 96%, 79% and 85% of cases respectively. However, in the other data-set-NAM combinations, snv and dt represented at least 25% of best performance and up to 42% in rice1_flour. Figure S4 represents predictive abilities for all classical-pre-processing methods for all traits in each data-set-NAM combination. For each data-set-NAM-trait combination, the inner loop of cross validation revealed either one outstanding pre-processing method, or two or three pre-processing methods that had similar predictive abilities. This explains why in general there is such variability in the choice of the best pre-processing as this choice is affected by small variations between repetitions of cross validation that are not likely to be really significant.

**Figure 5:**
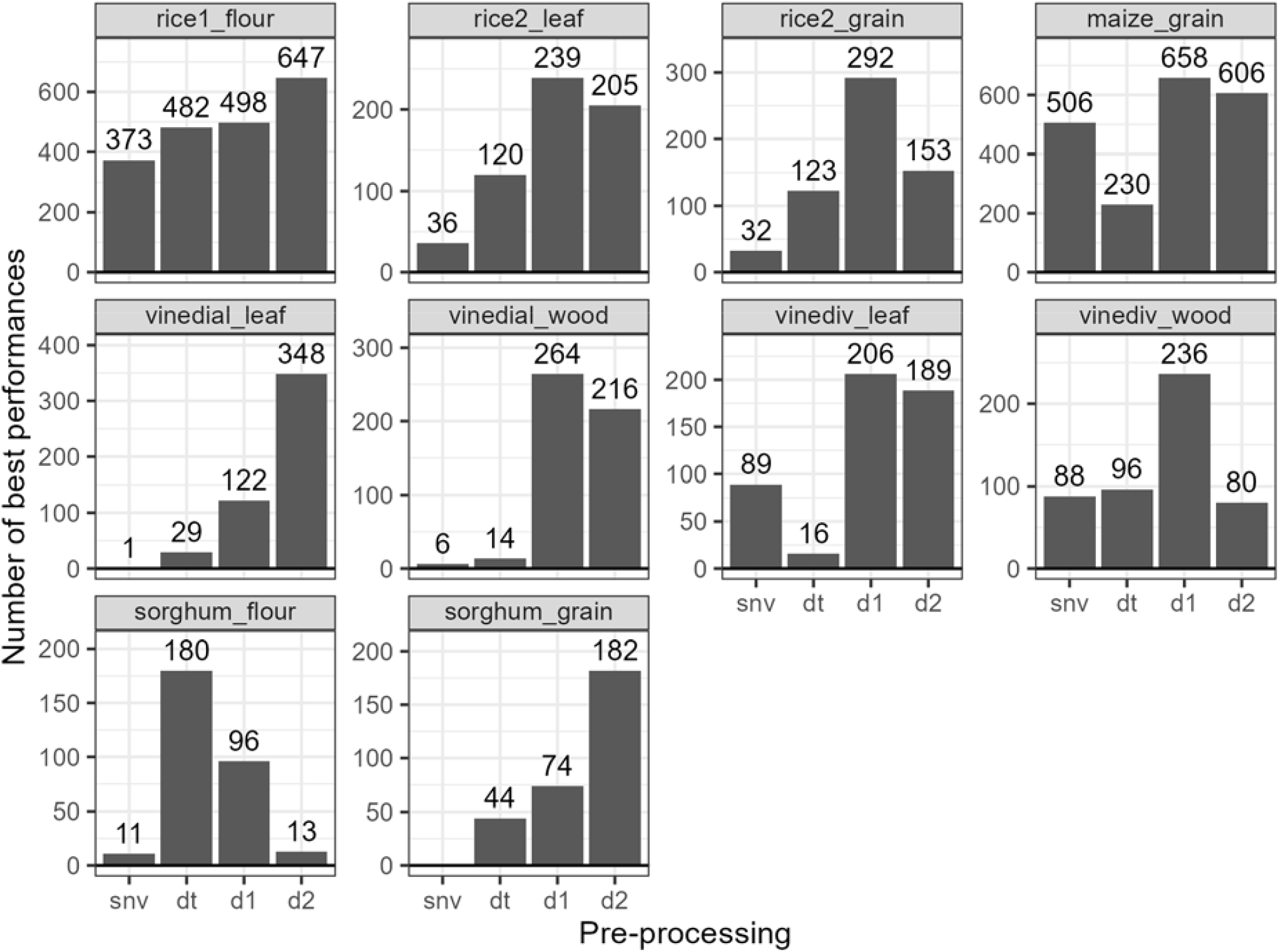
Number of times each pre-processing was the best performing. For each data set, each NIRS acquisition material (NAM) each trait and each cross-validation repetition, a different pre-processing method performed best in the inner tests on training data only. The number of times each pre-processing method performed best is summarized for each NAM in each data set. snv = standard normal variate, dt = detrend, d1 = first derivative after snv, d2 = second derivative after snv.

Figure 6 represents the number of times each number of PC was the best performing in each data-set-NAM combination across traits (according to the inner loop of cross validation). For sorghum_flour and sorghum_grain, a clear peak indicated that around 25 PC was the best performing selection. For vinedial_wood and maize_grain, peaks indicated that 30 and 75 PC respectively were the optimal number of PC, while for vinediv_leaf it was rather 10 PC. For other data sets, no clear signal could be detected, indicating that the optimal number of PC was highly variable. However, except for the vinediv and rice 2 data sets, overall between one and 10 PC were never or rarely optimal,indicating that selecting too few PC was generally disadvantageous.

**Figure 6:**
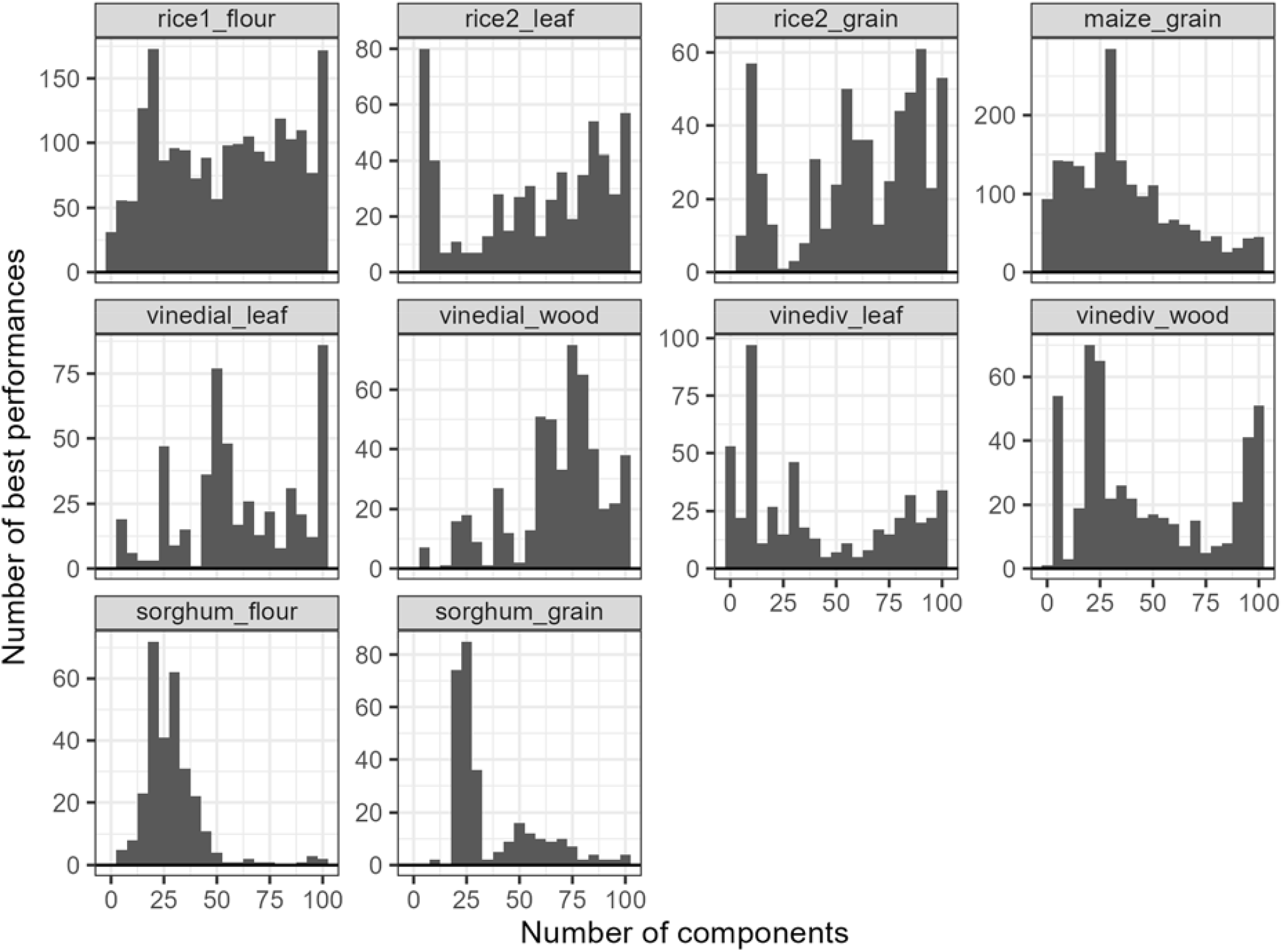
Histogram of the best performing number of components for the U method. For each data set, each NIRS acquisition material (NAM) each trait and each cross-validation repetition, a different number of components performed best for the U method in the inner tests on training data only. The number of times each number of components performed best is summarized for each NAM in each data set. Each bar represents an interval of 5 PC. snv = standard normal variate, dt = detrend, d1 = first derivative after snv, d2 = second derivative after snv.s

Figure S5 represents the median predictive ability reached by every number of PC for each trait in each data-set-NAM combination. These results showed that PA increased with the number of selected PC before reaching a plateau in most cases. This confirmed that selecting to few PC was disadvantageous, and explained the variability in the number of PC retained in the inner loop of cross validation. Similarly to what was observed between classical pre-processings, different number of components tended to yield very similar results. As a result, the choice between different number of PC was affected by small variations between repetitions of cross validation that do not reveal real differences between the number of PC.

For sorghum_flour, sorghum_grain, and maize_grain, PA did not reach a plateau but decreased after a certain number of PC, explaining why a clear peak could be observed for these data-set-NAM combinations. For vinediv and rice 2 data sets, there was either no increase of PA with the number of PC, or the plateau was reached when selecting less than five PC explaining that this data_set-NAM combination had a peak at such low numbers of PC.

## DISCUSSION

In this study, we investigated the performance of a SVD-based pre-processing of NIR spectra for PP using five data sets representing four species, and six type of traits with different genetic architectures. Two variants of the SVD were tested, either using scaled principal components, or using principal components weighted by their eigenvalues.

### Less parameter optimization for similar results

The SVD-based pre-processing method produced similar results to those of the best classical pre-processing method most of the time. Thus, our result show that the SVD-based pre-processing has the potential to become a go-to pre-processing as it gives high chances of performing as well as the best classical pre-processing methods, without having to test and choose among several pre-processing methods. Moreover, our result show that PA increase with the number of PC, and that most of the variation in PA occurred between one and 30 PC.

Consequently, if one does not have resources or time to optimize the number of PC, there are high chances that choosing any number of PC around or above 30 will lead to PA close to the optimum. It would be interesting to have a more in-depth study of the number of PC over which overfitting occurs and PA drops (which happened in some of our data sets).

Another advantage of working on PC instead of using classical pre-processing is the limited number of parameters that need to be optimized. The SVD-based method has only one parameter that could be optimized, namely the number of PC. On the other hand, the derivative methods which were the best classical pre-processing methods in most cases in this study have up to three parameters to optimize: derivative order, polynomial order and filter length for the Savitsky-Golay algorithm. Meyenberg et al. (2024) showed that it is possible to increase PA of PP by optimizing parameters of the Savitsky-Golay smoothing method. They found that the best parameter combination was dependent on the data set. Likewise, Braun et al. (2025) explored 677 combinations of parameter values of the Savitsky-Golay filter. They did not find any general pattern, or parameter combination that would increase PA for every trait or in every data set.

SVD or PCA based methods generally aim to reduce the dimensionality of data and several criteria exist to choose a number of components without having to optimize it with a nested cross validation scheme. In this study, we also tried to implement a classic PC selection criterion for PCA: keeping PC with eigenvalues superior to 1 (Kaiser, 1960). This method selected only one or two PC for each data set (results not shown) which would generally decrease PA according to our results. Moreover, it occurred that all PC had eigenvalues inferior to one, making this criterion not suitable for the analyses. Some authors have criticized this criterion because it is arbitrary, not supported by empirical evidences of its relevance, tends to over or under-extract PC, and failed to extract the right number of PC in simulated data in most cases (Morton & Altschul, 2019).

Even though other more robust methods for PC extraction exist, they are meant to reduce the dimensionality of the data, while the goal of the SVD-based pre-processing is to work on informative variables, reducing the dimensionality of spectra being a side advantage of it. Our results show that selecting less than ten PC generally yields lower PA than selecting more than 30 PC. This indicates that reducing dimensionality too much decreases the performance of PP and that relevant information is carried by PC that represent small amounts of spectral variance and would consequently typically be discarded by selection criteria. However, a criterion based on genetically relevant information could be useful. In this sense, it could be interesting to try and select PC linked to genetic parameters, *e*.*g*., selecting PC with the highest heritability.

### PC with small amount of information matter

One interesting outcome of this study is that the first few PC of raw spectra are generally worth more than 99% of the total variance of raw spectra (figure S6). Yet, selecting only these first few PC generally resulted in lower PA than selecting many more of the following PC, carrying all together less than 1% of spectral variance. This may show that in spectra, relevant information to discriminate genotypes is found in subtle variations that is represented in PC with small eigenvalues. This may also explain why the U pre-processing method performed better than the UD pre-processing method, as the U method gives more weight to such small components in the estimation of spectral genotypic BLUEs.

From a chemometrics point of view, it is not surprising that a small part of spectral variance could be relevant for prediction as chemometricians often predict chemical information that represents a small part of spectral variations while physical information is the most prominent information in spectra (Bull, 1991; Cowe & McNicol, 1985). The importance of PC representing small parts of the total variance has also been discussed in the PCR framework (Jolliffe, 1982; Næs & Martens, 1988), pointing out their importance in prediction.

### Why is working on PC as efficient as pre-processing spectra?

The present results show that working on PC of raw spectra instead of wavelengths can reach PA similar or sometimes higher than the ones achieved with the best classical pre-processing methods. Interpreting the meaning of selected components could help explaining the performance of the SVD-based pre-processing method. In fact, the first few PC carry most of the variance and are thus in the directions that scatters spectra the most. Such sub-spaces are more likely to bear the physical information of spectra as this is the information that causes the most variability in spectra (Bull, 1991; Cowe et al., 1985). Moreover, there are many examples in the chemometrics literature where the first PC of spectra is interpreted as representing particle size of milled samples, thus physical information (Cowe et al., 1985; Cowe & McNicol, 1985; Devaux et al., 1995; Otsuka, 2004; Otsuka et al., 2003; Rantanen et al., 2005). In a review article, Caporaso et al. (2018) reported that first PC of spectra on whole grains of wheat, barley and sorghum were related to physical effects such as non-uniformity of illumination, grain shape and surface texture. Thus, selecting only the first few PC is likely to select mostly physical information. Conversely, discarding these PC would likely select chemical information while removing physical information, and having an effect similar to classical pre-processing.

Selecting many components while scaling them as in the U pre-processing gives the same importance the first components with high eigenvalues and to components with low eigenvalues. Thus, it probably enhances the importance of the chemical signal in spectra and lowers the weight of physical information when computing genetic value of spectra. On the other hand, classical pre-processing methods are meant to remove the physical part of spectral information. Thus, the U method and classical pre-processing methods probably have similar effects on spectra. However, unlike classical pre-processing methods, the U method does not completely remove physical effects, but only gives more weight to chemical information. Put together, our results may show that the chemical information contained in spectra is the most relevant for PP (classical pre-processing performs better than raw spectra), but that physical information can also be valuable (SVD-based method performs similarly to classical methods and sometimes outperforms them).

The performances of the SVD-based method may also partly be explained by the features that make PCR useful when working on spectra, namely, handling multicollinearity and working on orthogonal informative, less noisy PC (Jolliffe, 1986). Indeed, estimating the genetic part of spectra on such PC likely produces better estimation than doing it on many correlated, noisy, less informative wavelengths. Working on more informative subsets of spectra for PP is analogous to working with more informative subset of markers (using functional annotation or other information about markers) in GP, which has been reported to improve its PA (Campos et al., 2013; Edwards et al., 2016; Gao et al., 2015, 2017; MacLeod et al., 2016; Mollandin et al., 2022; Ramstein et al., 2016; Speed & Balding, 2014).

### Multivariate methods in prediction

Whether it is GP or PP, high dimensional datasets are used to predict genetic values. In both cases, large sets of variables are mobilized (wavelengths or molecular markers), some of theme carrying similar information (linkage disequilibrium, correlation between wavelengths) some of them known to be more useful (functional annotations), some of them carrying non additive information (epistasis, GxE interaction) and others for which their interest is still unknown. In both cases, multivariate methods that sum up these large sets into fewer, more informative and independent variables could be used to ease computation and help finding useful subspaces of the predictor data. As mentioned in the introduction, the SVD-based pre-processing relates to methods such as PCR or PLSR. However, it differs from such methods as PCR and PLSR use subspaces of explanatory variables to model one or multiple response variables. The SVD-based pre-processing is not a model, and it uses explanatory variables to model subspaces of the response variable which are meaningful compared to the original response variables.

Our results show that it is better to estimate the genetic part of spectra on PC instead of wavelengths. Thus, one could argue that PLSR is in theory well suited for prediction of genetic values using high dimensional data. Indeed, it finds sub-spaces of the predictors that correlate the best to the data or to sub-spaces of the data to predict. It has been tested in several PP studies with results always in the same range of PA as the reference prediction method used (Adunola et al., 2024; Bienvenu et al., 2025; Dallinger et al., 2023; Gonçalves et al., 2021; Lane et al., 2020; Meyenberg et al., 2024), except for Gonçalves et al. (2021) where BayesB model outperformed PLSR. Classical PLSR attributes weights to the latent variables as in PCA. Our results show that working on scaled PC is important for the SVD-based pre-processing to work. Thus, it would be interesting to test a PLSR method with scaled latent variables. PLSR has also been tested in a few GP studies, showing that it can have lower PA than the reference method (Solberg et al., 2009), similar PA (Singh et al., 2016), or better PA in the context of multi-environment-trait or GxE predictions (Montesinos-López et al., 2022a, 2022b; Ortiz et al., 2023). Such results are encouraging and it would be interesting to try multi-block PLSR to gather genomic and phenomic information in multi-environment models. There are also a few instances of GP using the PCR method (Dadousis et al., 2014; Du et al., 2018; Macciotta et al., 2010; Pintus et al., 2012; Solberg et al., 2009). PA were generally slightly lower (much lower in Solberg et al. (2009)) with PCR than with classical GP models. Poorest performance of PC-based methods in genomic could be explained by the lower redundancy and multicollinearity of marker data compared to NIR spectra.

Studies reporting slightly lower PA for PCR based GP also reported substantial computational efficiency gains compared to GBLUP models. In the present study, computational efficiency gains were achieved in the spectra pre-processing step but not in the prediction step, and are more a side effect of using PCA rather than an initial goal. However, it can become a major stake for sustainable research. Data centers and computational facilities have a water and carbon footprints large enough for the scientific community to try to measure and reduce it (Jiang et al., 2025; Lannelongue et al., 2023; Siddik et al., 2021), even though scientific computation is not the only use of data centers and computing facilities. Recent advances and wide deployment of artificial intelligence (AI) is likely to increase the environmental impacts of computing facilities, and controversy about the use of resources for AI is ongoing (Nordgren, 2022). Thus, it is relevant to try to find a balance between computational efficiency and model’s performance. It is especially true for PP, developed to be a cost efficient, high throughput genetic value prediction method, that could be used with little financial means, or for species that cannot be routinely genotyped yet. As PP can be advertised as a low-tech prediction method, computational efficiency could become an important feature to consider when studying it.

## CONCLUSION AND PERSPECTIVES

In this study, we investigated the performance of a SVD-based pre-processing of NIR spectra for PP using five data sets representing four species, and six type of traits with different genetic architecture. We showed that this pre-processing performs in most cases as well as the best classical chemometric pre-processing among the ones tested, making it a possible go-to pre-processing. Moreover, it requires less parameter optimization and reduces the computation time of spectral genotypic BLUEs. Exploring different threshold for component selection revealed that the most relevant information for PP is contained in components carrying small parts of spectral variance. Thus, it is likely that the chemical information contained in spectra drives PP performance even though physical information remains valuable.

A wider use of multivariate chemometrics methods could benefit PP. Such methods can help derive meaningful variables from spectra. Thus, they could open prospects on understanding PP better and verify if it can better predict non additive information such as GxE or GxG interactions.

## Abbreviations

GP: genomic prediction
GW: grain/fruit weight related traits
MORPH: morphology related traits of spike/ear/panicle/grape
NAM: Near infrared spectroscopy acquisition material
PH: plant height
PA: predictive ability
PC: principal component
PCA: principal component analysis
PCR: principal component regression
PHENO: phenology related traits
PLSR: partial least square regression
PP: phenomic prediction
VIG: vigour related traits
YIELD: yield related traits

## AUTHOR CONTRIBUTIONS

**CB:** Conceptualization; data curation; formal analysis; investigation; methodology; writing— original draft. JMR: Methodology. **M. Séne:** data curation; writing – original draft. **SACP:** writing – original draft, data curation. **M. Singer:** data curation. **BLF:** data curation. **NT:** funding acquisition; project management; writing—review and editing. **FDB:** data curation. **DP:** Conceptualization; funding acquisition; investigation; supervision; validation; writing—review and editing. **HDV:** Conceptualization; funding acquisition; investigation; supervision; validation; writing—review and editing. **VS:** Conceptualization; funding acquisition; investigation; supervision; validation; writing—review and editing.

## ACKNOWLEDGMENTS

The work of Clément Bienvenu was supported by a doctoral allowance from the French Ministry of Higher Education and Research. Mamadou Séne received support for his PhD grant from CIRAD and INRAE CLIMAE Metaprogramme. The sorghum dataset was produced in the frame of the NITROSORG project (2022-2025) that has been funded by CASDAR Semences et Sélection Végétale (AAP 2021 C2021-06). The rice dataset 2 was produced within the framework of the DINAAMICC project (contract FOOD/2021/422-791), funded by the European Union under the DeSIRA program. Authors would like to thank Patrice Jeanson and Clothilde Boubée de Gramont from LIDEA seeds / Eurosorgho, Joël Alcouffe, Quentin Duprat and Philippe Dufour from RAGT2N, and Florentin Pietrus from CIRAD for their help in setting up the trials for the NitroSorg project.

## DATA AVAILABILITY STATEMENT

All data used in this study are available online:

- Rice 1: All files are available from the FigShare database (https://doi.org/10.6084/m9.figshare.27282459.v1).
- Rice2: Currently being published on the cirad dataverse but no permalink available yet
- Maize: All files are available on GitHub (https://github.com/ajdesalvio/Maize-NIRS-GBS
- Grapevine: All files are available at the INRAE data portal (spectra available at https://doi.org/10.15454/BICRFX, phenotypic data for the diversity panel are available at https://doi.org/10.15454/8DHKGL, phenotypic data for the half diallel populaion are available at https://doi.org/10.15454/PNQQUQ)
- Sorghum: Currently being published on the cirad dataverse but no permalink available yet

Codes are available on Gitlab : https://gitlab.cirad.fr/agap/giv/phenomic_chemometric_preprocessing/-/tree/696c376e81c7296bf1cf1b52b5356767a44baccf/

## SUPPLEMENTAL MATERIAL

**Table S1: Mean and standard deviations of traits for the sorghum and rice data set 2**.

**Figure S1: Variance decomposition for traits of the sorghum data set**. GW = grain weight or berry weight, PH = plant height, PHENO = phenology. Numbers in black are heritabilities of each trait.

**Figure S2: Variance decomposition for traits of the rice data set 2**. GW = grain weight or berry weight, PH = plant height, PHENO = phenology, MORPH = morphology of the grain/fruit bearing organ, VIG = vigour. Numbers in black are heritabilities of each trait.

**Figure S3: Comparison between predictive abilities obtained with raw spectra, the best classical chemometric method, the U method and the combination of U and chemometric methods**. In the RAW method, genotypic BLUEs were estimated on wavelengths of raw spectra. In the CHEMO method, genotypic BLUEs were estimated on the wavelengths of pre-processed spectra. In the U method, genetic values were estimated on the U matrix of a singular value decomposition of raw spectra. In the CHEMO + U method, genetic values were estimated on the U matrix of a singular value decomposition of classically pre-processed spectra. For each cross-validation repetition, trait, NIRS acquisition material (NAM) in each data set, the number of components kept, the best performing classical pre-processing, and the best combination of the two were determined by an inner five-fold cross validation on the training data. GW = grain weight or berry weight, PH = plant height, PHENO = phenology, MORPH = morphology of the grain/fruit bearing organ, VIG = vigour.

**Figure S4: Influence of classical pre-processing methods on the predictive abilities in the inner cross validation loop**. GW = grain weight or berry weight, PH = plant height, PHENO = phenology, MORPH = morphology of the grain/fruit bearing organ, VIG = vigour. snv = standard normal variate, dt = detrend, d1 = first derivative after snv, d2 = second derivative after snv.

**Figure S5: Evolution of predictive abilities with the number of components chosen in the inner cross validation loop**. Each point represents de median predictive ability for a given number of components in the inner loop. GW = grain weight or berry weight, PH = plant height, PHENO = phenology, MORPH = morphology of the grain/fruit bearing organ, VIG = vigour.

**Figure S6: Scree plot of the principal components of the singular value decomposition of spectra**. Singular value decomposition were done on raw spectra and using all available spectra without partitioning into training and validation data.

