## Supplementary materials for "OPTIMIZING PRE-PROCESSING OF NEAR INFRARED SPECTRA FOR PHENOMIC PREDICTION USING SINGULAR VALUE DECOMPOSITION"

6: Dispositif en Partenariat Système de Production d’Altitudes Durable (DP-SPAD), Antsirabe, Madagascar

7: Geno-Vigne, IFV-INRAE-Institut Agro, F-34398, Montpellier, France

Vincent Segura, Hugues de Verdal, and David Pot should be considered joint senior authors.

6 figures, 1 table

***Table S1: Mean and standard deviations of traits for the sorghum and rice data set 2.***

|  | Sorghum_mond_2023 | | Sorghum_riv_2023 | | Sorghum_riv_2024 | | Rice2_and_fm | | Rice2_and_fu | | Rice2_tal | |
| --- | --- | --- | --- | --- | --- | --- | --- | --- | --- | --- | --- | --- |
|  | mean | sd | mean | sd | mean | sd | mean | sd | mean | sd | mean | sd |
| PHENO | \ | \ | 88.8 | 9.9 | 80.2 | 10.9 | 45731 | 5.4 | 45731 | 3.7 | 45707 | 4.1 |
| PH | 186.5 | 72 | 201.6 | 92.6 | 179.9 | 74.6 | 98.4 | 10.6 | 77.7 | 8.7 | 110.9 | 12.2 |
| GW | 23.3 | 6.3 | 22.9 | 6.8 | 26 | 6.2 | 5.5 | 0.72 | 5.3 | 0.62 | 5.3 | 0.49 |
| VIG | \ | \ | \ | \ | \ | \ | 12.5 | 2.8 | 11 | 3.6 | 77.1 | 19.3 |
| MORPH | \ | \ | \ | \ | \ | \ | 19.4 | 1.7 | 18.9 | 1.5 | 21.44 | 1.9 |
| YIELD | \ | \ | \ | \ | \ | \ | 705.1 | 206.1 | 279.9 | 167.5 | 602.5 | 189.2 |

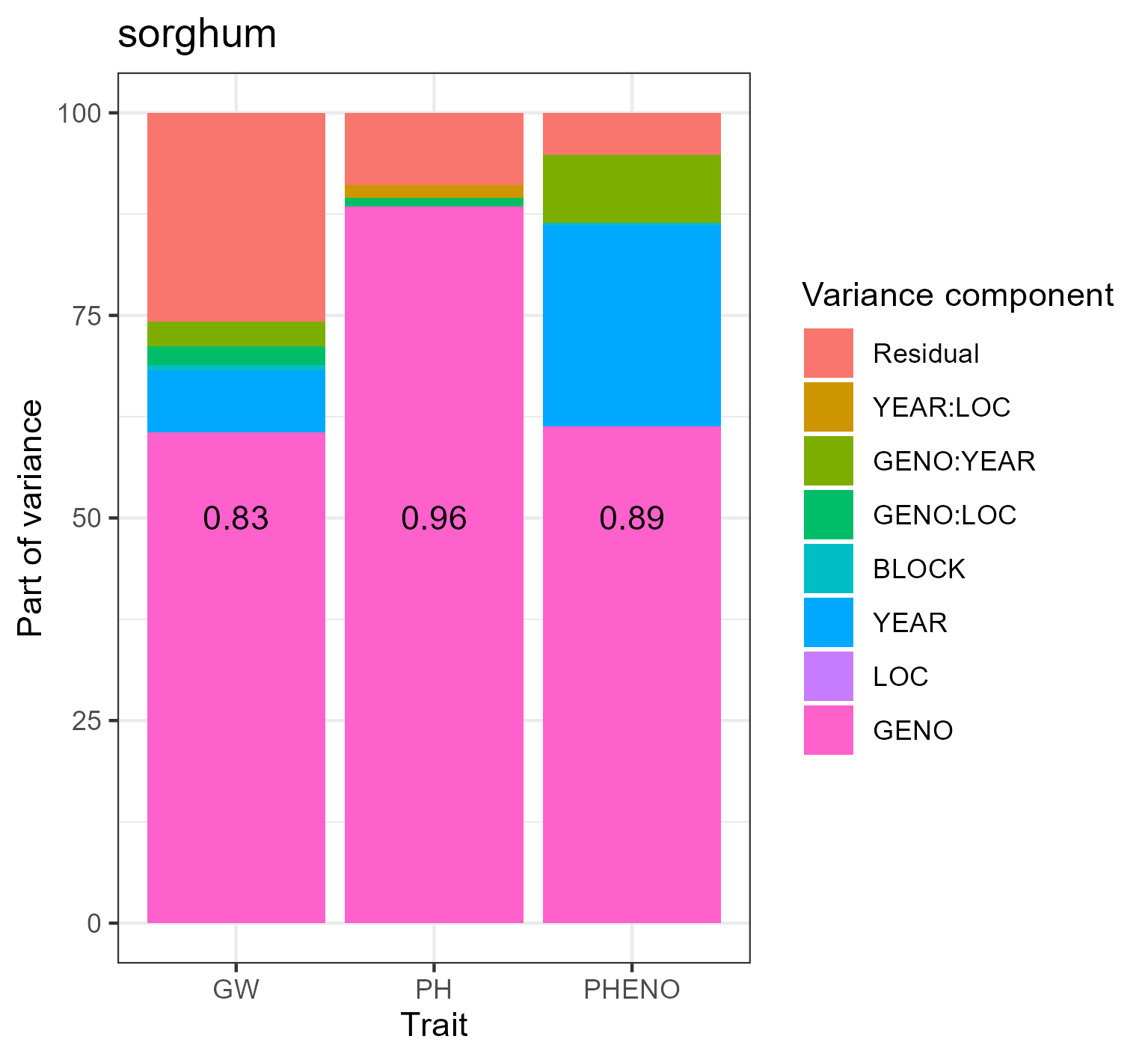

***Figure S1: Variance decomposition for traits of the sorghum data set****. GW = grain weight or berry weight, PH = plant height, PHENO = phenology. Numbers in black are heritabilities of each trait.*

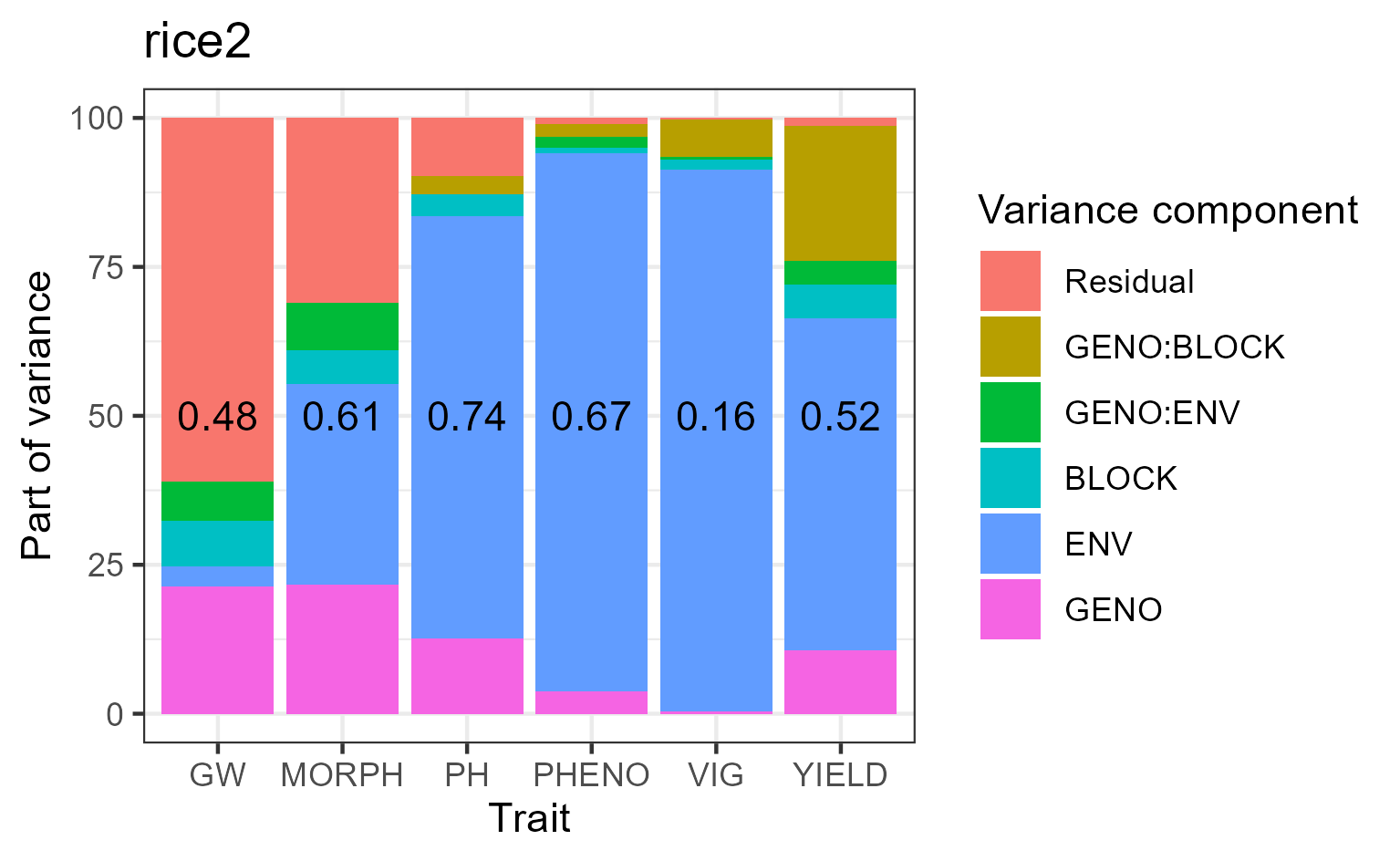

***Figure S2: Variance decomposition for traits of the rice data set 2****. GW = grain weight or berry weight, PH = plant height, PHENO = phenology, MORPH = morphology of the grain/fruit bearing organ, VIG = vigour. Numbers in black are heritabilities of each trait.*

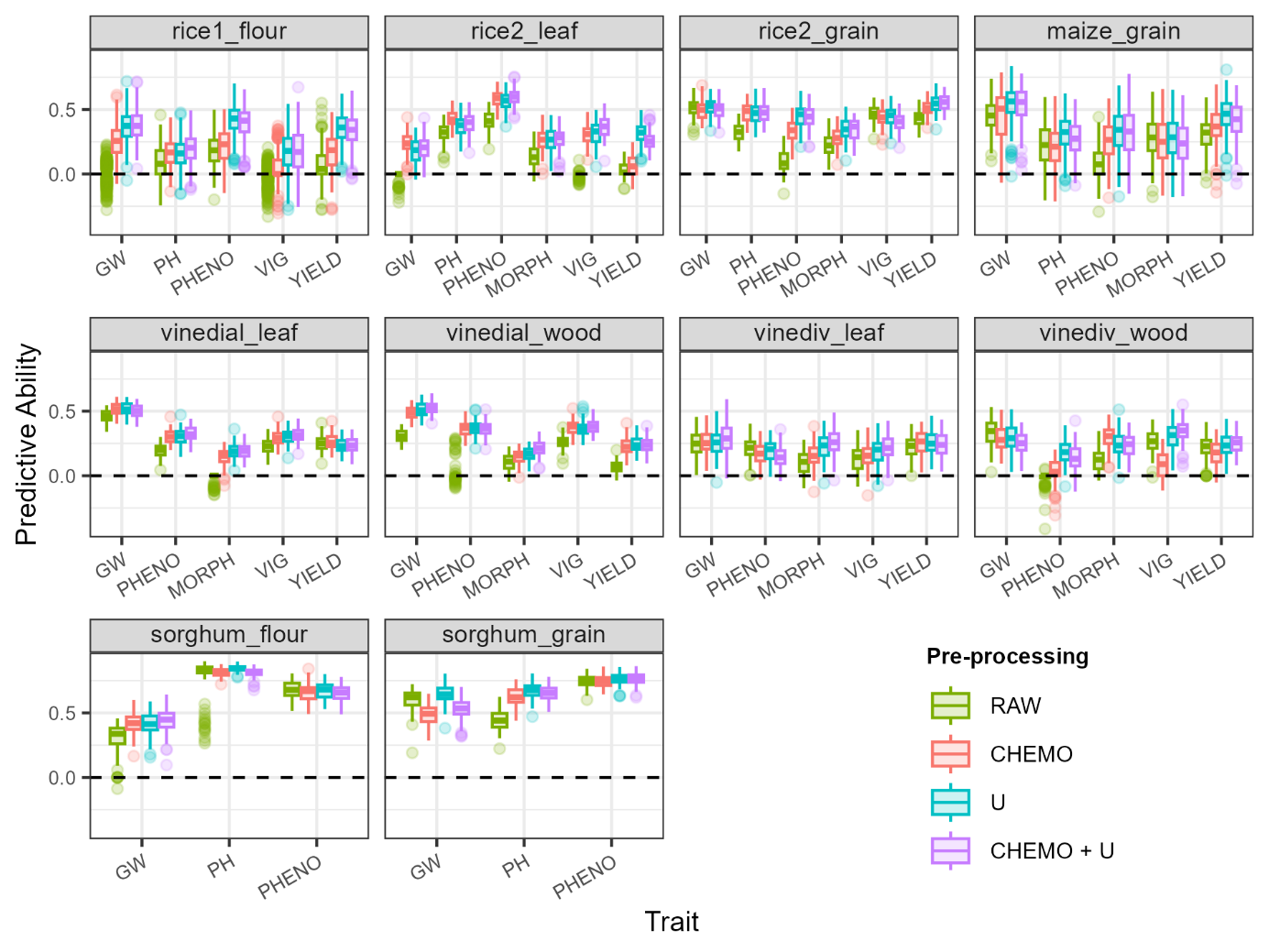

***Figure S3: Comparison between predictive abilities obtained with raw spectra, the best classical chemometric method, the U method and the combination of U and chemometric methods.*** *In the RAW method, genotypic BLUEs were estimated on wavelengths of raw spectra. In the CHEMO method, genotypic BLUEs were estimated on the wavelengths of pre-processed spectra. In the U method, genetic values were estimated on the U matrix of a singular value decomposition of raw spectra. In the CHEMO + U method, genetic values were estimated on the U matrix of a singular value decomposition of classically pre-processed spectra. For each cross-validation repetition, trait, NIRS acquisition material (NAM) in each data set, the number of components kept, the best performing classical pre-processing, and the best combination of the two were determined by an inner five-fold cross validation on the training data. GW = grain weight or berry weight, PH = plant height, PHENO = phenology, MORPH = morphology of the grain/fruit bearing organ, VIG = vigour.*

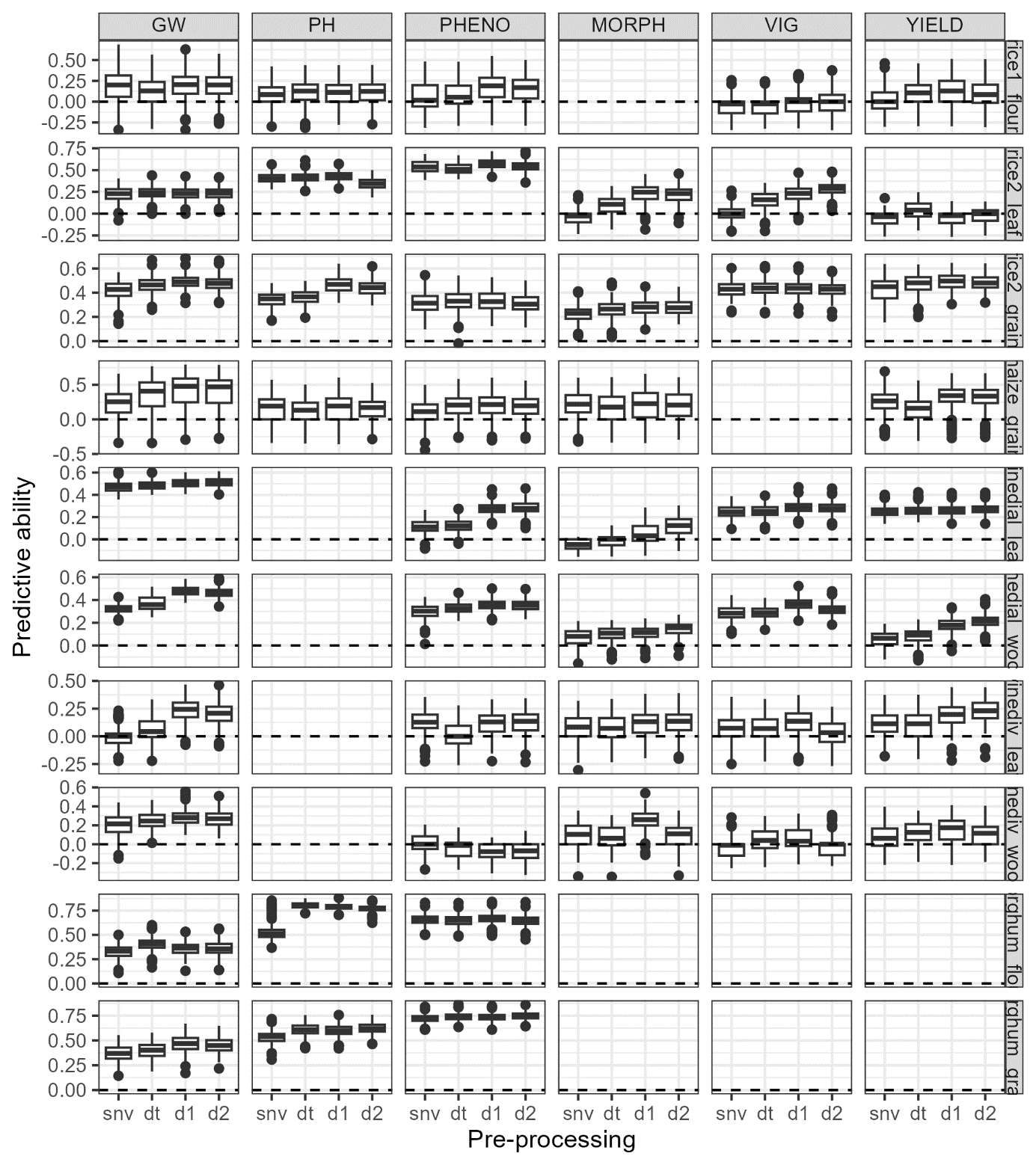

***Figure S4: Influence of classical pre-processing methods on the predictive abilities in the inner cross validation loop.*** *GW = grain weight or berry weight, PH = plant height, PHENO = phenology, MORPH = morphology of the grain/fruit bearing organ, VIG = vigour. snv = standard normal variate, dt = detrend, d1 = first derivative after snv, d2 = second derivative after snv.*

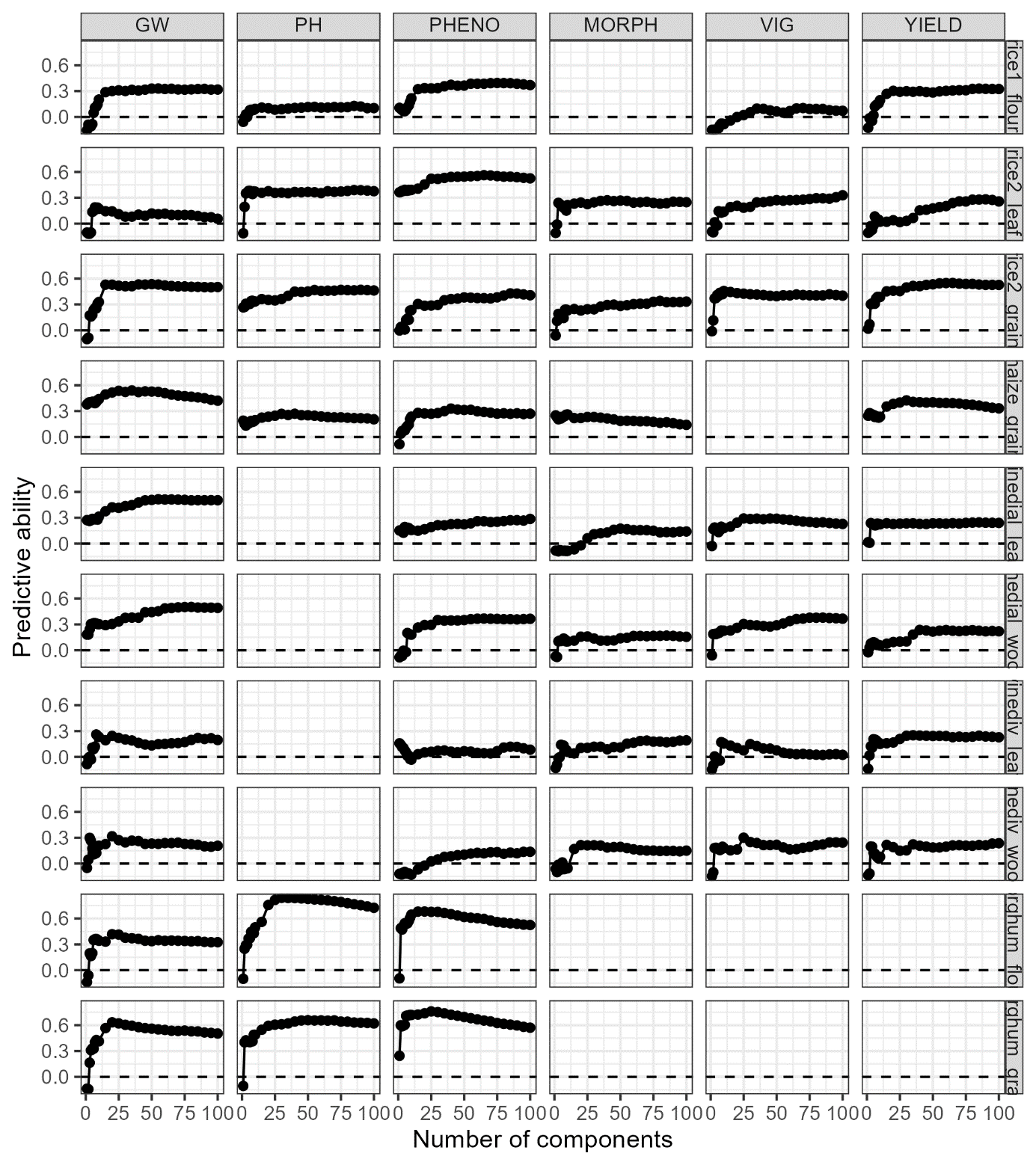

***Figure S5: Evolution of predictive abilities with the number of components chosen in the inner cross validation loop.*** *Each point represents de median predictive ability for a given number of components in the inner loop. GW = grain weight or berry weight, PH = plant height, PHENO = phenology, MORPH = morphology of the grain/fruit bearing organ, VIG = vigour.*

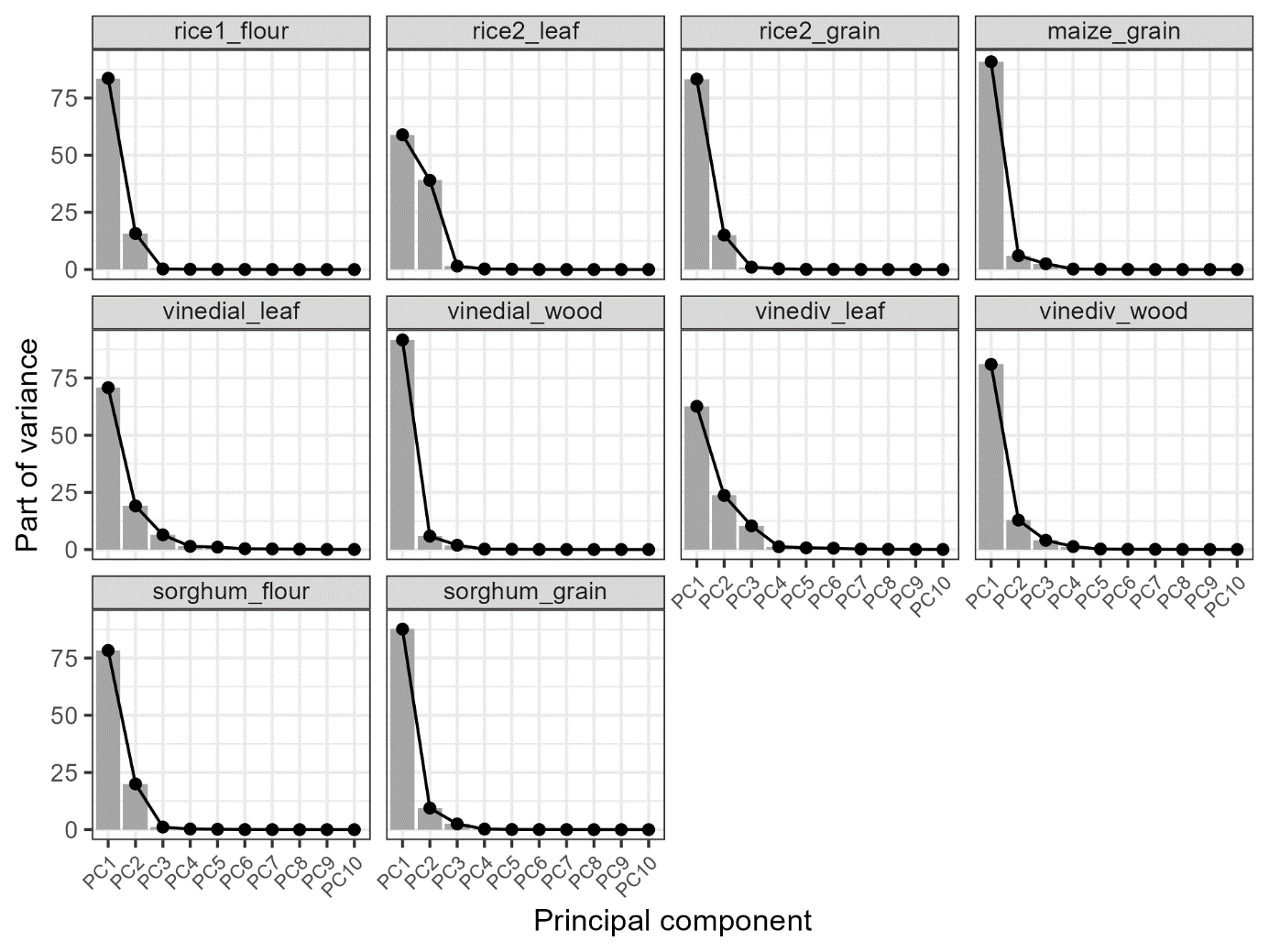

***Figure S6: Scree plot of the principal components of the singular value decomposition of spectra.*** *Singular value decomposition were done on raw spectra and using all available spectra without partitioning into training and validation data.*
